# Negative and neutral interactions are prevalent in interactions between marine chitin degraders

**DOI:** 10.64898/2026.06.17.732797

**Authors:** Vilhelmiina Haavisto, Peter F. Doubleday, Andreas Sichert, Uwe Sauer

**Affiliations:** Institute of Molecular Systems Biology, ETH Zürich, Zürich, Switzerland; Life Science Zurich PhD Program on Microbiology and Immunology, Zurich, Switzerland

**Keywords:** Chitin degradation, marine bacteria, polysaccharide degradation, interactions, inhibition, marine snow

## Abstract

Bacterial chitin degradation contributes to global carbon cycling, particularly in marine environments where it is a highly abundant polysaccharide. Despite the taxonomic diversity of co-occurring chitin-degrading bacteria, the influence of individual traits on interactions between them remains poorly understood. Here, we measured key physiological traits of seven chitin degraders and investigated how these traits shape interaction outcomes and chitin degradation in pairwise cocultures. We found mainly negative and neutral interactions among degraders, contrasting with the synergistic dynamics observed with other complex polysaccharides. However, chitin degradation was not consistently diminished. These interaction types could be attributed to the limited partitioning of degradation products, alongside variations in enzyme repertoires and attachment behaviours that help some degraders to prevail over others. Further, we showed that one degrader can strongly inhibit the growth of others, even those possessing favourable physiological traits, likely due to the secretion of inhibitory compounds. These findings extend our understanding of the breadth of interactions among primary polysaccharide degraders and their implications for the degradation process.

**One-sentence Summary:** The physiological traits of bacteria that degrade chitin, a highly abundant biopolymer in marine environments, promote a range of neutral and negative interactions among them.

## Introduction

Microorganisms contribute to elemental cycles by recycling organic matter across diverse environments including soil, animal digestive tracts, and the marine environment (Falkowski, Fenchel, and Delong 2008). In the ocean, heterotrophic bacteria remineralize chemically diverse particulate organic matter (POM), also known as marine snow (Azam *et al*. 1983, Smith *et al*. 1992). One component of POM is chitin, one of the most abundant marine polysaccharides whose annual production exceeds 1 billion tons (Souza *et al*. 2011). Chitin consists of *β*-1,4-linked chains of *N*-acetylglucosamine (GlcNAc) monomers. The antiparallel chain arrangement in α-chitin, the most abundant form, renders it highly crystalline and insoluble, though not resistant to degradation (Souza *et al*. 2011). Bacteria that express extracellular and membrane-bound enzymes (broadly known as chitinases) are able to degrade chitin into chitooligomers of different lengths (Souza *et al*. 2011, Beier and Bertilsson 2013). These chitooligomers can be imported into the peri- and cytoplasm by various transport systems (Eisenbeis *et al*. 2008, Hunt *et al*. 2008, Monge and Gardner 2021), where GlcNAc is deacetylated and deaminated to produce fructose-6-phosphate for entry into central metabolism (Itoh and Kimoto 2019).

The degradation of chitin and other components of POM by heterotrophic bacteria significantly contributes to the overall flux and fate of carbon in the ocean (Nguyen *et al*. 2022, Iversen 2023). Degradation is modulated by a combination of abiotic and biotic factors, including enzyme secretion (Arnosti 2011), substrate attachment (Kiørboe *et al*. 2003, Ebrahimi, Schwartzman, and Cordero 2019), and interactions with other bacteria (Beier and Bertilsson 2013, Daniels *et al*. 2022, D’Souza *et al*. 2023). Extracellular chitin degradation creates a public resource pool that supports multispecies particle-associated communities, including not only chitin degraders but also a range of metabolically diverse, non-degrading bacteria (Datta *et al*. 2016, Szabo *et al*. 2022). While interactions between degraders and non-degraders have been studied (Pontrelli *et al*. 2022, 2024, Daniels, Vliet van, and Ackermann 2023), interactions among degraders and their implications for chitin degradation are not well understood. This limits our understanding of how chitin degradation proceeds in marine environments, where interactions among taxonomically diverse, co-occuring degraders on the microscale could impact the fate of carbon in sinking POM, and thus global marine carbon cycling (Nguyen *et al*. 2022).

The degradation of complex substrates can facilitate various types of interactions, ranging from competition (Shi, Odt, and Weimer 1997, Chen J and Weimer 2001) to the evolution of cheating (Jagmann, Styp von Rekowski, and Philipp 2012, Enke *et al*. 2018) or cooperation through division of labour (Rakoff-Nahoum, Foster, and Comstock 2016, Lindemann 2020). Interactions such as exploitation between chitin degraders and non-degrading strains that consume GlcNAc have been shown experimentally (Pollak *et al*. 2021, Pontrelli *et al*. 2022). However, whether similar dynamics exist among degrading strains is only known for one degrader pair (Jagmann, Styp von Rekowski, and Philipp 2012). Interaction outcomes depend on several factors, including enzyme complementarity (Gálvez *et al*. 2020, Lindemann 2020) and resource niche partitioning between degraders (Lindemann 2020, Brochet *et al*. 2021). Although synergistic interactions between different chitin-degrading enzymes have been modelled (Guseva *et al*. 2024) and shown experimentally in single strains (Horn *et al*. 2006, Monge *et al*. 2018, Zhang *et al*. 2023), to our knowledge, cooperative chitin degradation by two or more strains has not been shown experimentally. Indeed, it has been suggested that chitin degradation is inherently non-cooperative (Enke *et al*. 2018).

To investigate how physiological traits influence the outcome of interactions between chitin degraders, we examined seven chitin-degrading strains that were previously isolated from a coastal marine environment using a particle-enrichment strategy (Datta *et al*. 2016, Enke *et al*. 2018). These strains have all previously been described as chitin degraders (Enke *et al*. 2018, Pollak *et al*. 2021, Pontrelli *et al*. 2022), and represent four genera across two orders **(Table 1)**. We began by assessing encoded and expressed enzyme repertoires during degradation of colloidal chitin through genomic analysis and extracellular proteomics. Next, we investigated the potential for niche partitioning of chitin-derived resources, allowing us to formulate hypotheses about how degraders may interact. To test these hypotheses, we then conducted pairwise co-culture experiments with a subset of the degraders. We also followed up a strong inhibitory interaction that we observed in some cocultures, and showed that it is likely driven by the production of an inhibitory compound by one of the degraders. Altogether, we aimed to discover the types of interactions occurring between chitin-degrading marine strains to shed light on how they may affect carbon cycling in marine environments.

**Table 1:**
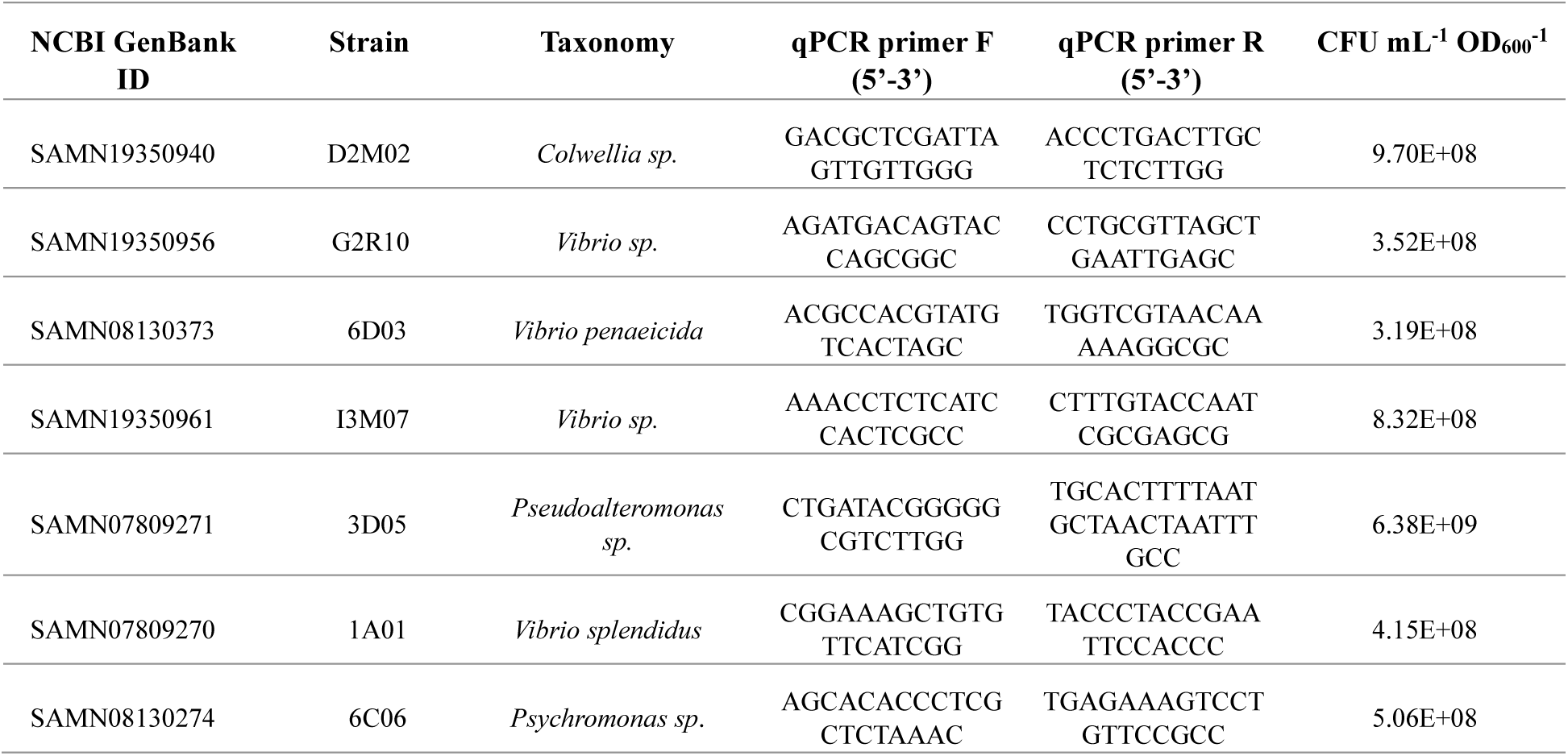
Chitin degrader strains used in this work alongside strain-specific qPCR primer sequences and CFU mL^-1^ OD_600_ ^-1^ values of each species grown in Marine Broth 2216 used to determine how much preculture is required to inoculate cultures at a given cell density.

## Materials and methods

### Growth media

All experiments were conducted with minimal marine media (hereafter MBL) (Pollak *et al*. 2021), with chemicals from Sigma unless otherwise noted. MBL was mixed fresh for experiments from stocks of its components: 1’000x concentrated trace minerals (FeSO_4_*7H_2_O, 2.1 g L^-1^; H_3_BO_3_, 30 mg L^-1^; MnCl_2_*4H_2_O, 100 mg L^-1^; CoCl_2_*6H_2_O, 190 mg L^-1^; NiCl_2_*6H_2_O, 24 mg L^-1^; CuCl_2_*2H_2_O, 2 mg L^-1^; ZnSO_4_*7H_2_O, 144 mg L^-1^; Na_2_MoO_4_*2H_2_O, 36 mg L^-1^; NaVO_3_, 25 mg L^-1^; NaWO_4_*2H_2_O, 25 mg L^-1^; Na_2_SeO_3_*5H_2_O, 6 mg L^-1^), sodium phosphate dibasic (1 mM), sodium sulfate (1 mM), TES buffer (1M, pH 8.2), 4x concentrated seawater salts (NaCl, 80 g L^-1^; MgCl_2_*6H_2_O, 12 g L^-1^; CaCl_2_*2H_2_O, 0.6 g L^-1^; KCl, 2 g L^-1^), autoclaved Milli-Q water, and a carbon source.

Colloidal chitin was made from chitin from shrimp shells (Sigma) by mixing 10 g chitin with 100 mL phosphoric acid (85% w/v). The mixture was incubated at 4°C for 48 hours and mixed with approximately 500 mL water and shaken vigorously to resuspend the chitin-acid slurry. The slurry was vacuum-filtered through filtration paper (Macherey-Nagel, MN615) to remove liquid. The resulting slurry was then placed into 3 kDa dialysis tubing cellulose membrane (Sigma-Aldrich) and dialysed in distilled water for 5-7 days, changing the water daily, to remove residual monomers, oligomers, and acid. The colloidal chitin was then collected from inside the tubing, homogenized using a blender, adjusted to pH 7 with NaOH, diluted to a concentration of 10 g L^-1^, and autoclaved before use.

For experiments with carbon sources other than colloidal chitin, the carbon sources were prepared as sterile 0.4 M stock solutions and stored at 4°C. The stock solution was added to freshly prepared MBL at a final concentration of 20 mM (GlcNAc) or 10 mM (acetate). For carbon sources other than chitin and GlcNAc, ammonium chloride was added as a nitrogen source at a final concentration of 10 mM.

### Strains

All strains were previously obtained from the laboratory of Otto Cordero at the Massachusetts Institute of Technology. The strains were originally isolated from coastal seawater at Canoe Beach, Nahant, MA, USA; 42°25’11.5“ N, 70°54’26.0” W (Datta *et al*. 2016, Enke *et al*. 2019). All strains were maintained as separate glycerol stocks stored at -70°C and streaked onto Marine Broth 2216 (Difco) 1.5% agar plates for use in experiments. The plates were stored at 4°C for at most 1 month. All growth experiments were carried out at 27°C with 200 RPM shaking in 24-deep-well plates unless otherwise specified.

### Chitin growth and quantification of chitin degradation

Growth on chitin was assessed by inoculating single colonies of each strain into Marine Broth 2216 in 15mL Greiner tubes and inoculating these overnight cultures into colloidal chitin MBL for a chitin preculture of between three and seven days. Upon reaching late-exponential phase, cultures were transferred onto fresh colloidal chitin for the growth assay. OD_600_ was measured daily, with denser sampling as cultures entered the exponential growth phase. Samples to quantify remaining chitin in the cultures were taken daily and stored at -20°C until processing and analysis. OD_600_ measurement was carried out as follows for this experiment and all other experiments involving growth on chitin. Plates were centrifuged for 1 minute at 500 RCF to lightly sediment the chitin while allowing bacterial cells to remain planktonic. 100 µL of culture was carefully removed and used to measure OD_600_ using a plate reader (Tecan Sunrise), then discarded. Chitin degradation was quantified according to a protocol based on degradation with an enzyme cocktail and quantification using a 3,5-dinitrosalicylic acid-based reagent (Pontrelli and Sauer 2025). A Spearman correlation was performed between growth and degradation rates using triplicate measurements from all degraders, as the data were not normally distributed.

### Growth assays on GlcNAc and acetate

Degrader strains were precultured in Marine Broth 2216 in 15 mL Greiner tubes. For the GlcNAc growth assay, all strains were first grown individually in a GlcNAc preculture before transfer into fresh GlcNAc MBL for the growth assay. For the acetate growth assay, cells from Marine Broth 2216 precultures were washed using MBL without a carbon source and inoculated into acetate MBL at a concentration of 2×10^6^ CFU mL^-1^. For the GlcNAc assay, OD_600_ was measured hourly to capture the exponential growth phase, whereas for acetate it was measured twice daily due to slower growth.

### Attachment assay

Degrader strains were precultured individually on Marine Broth 2216 in 15mL Greiner tubes and inoculated in quadruplicates into either MBL without a carbon source, or MBL with 2 g L^-1^ colloidal chitin, at a concentration of 3×10^7^ CFU mL^-1^. The cultures were incubated at 27°C with 200 RPM shaking for 10 hours, at which point they were centrifuged at 500 RCF for 1 minute to sediment the chitin. The planktonic phase was sampled for qPCR. The percentage of cells bound to chitin particles for each strain was calculated as a percentage of the total population (measured using DNA concentration as a proxy, see section ‘Strain-specific qPCR’), assuming an equal population size in chitin and no-chitin cultures.

### Enzyme repertoire

All degrader genomes were obtained from NCBI **(Table 1)**. To explore their CAZyme content, dbCAN 3 (Zheng *et al*. 2023) was used to annotate the genomes. Sequences of putative chitinases (GH18 and GH19 families) were extracted and the architecture of their catalytic domains was visualized using EMBL SMART (Letunic and Bork 2018, Letunic, Khedkar, and Bork 2021). A chitinase cladogram was constructed with the catalytic domain sequences using MEGA 11 (Tamura *et al*. 2011), using MUSCLE alignment for the protein sequences and a WAG + F + G maximum likelihood method to build the tree. SignalP 6.0 (Teufel *et al*. 2022) was used to predict signal peptide sequences within GH18 and GH19 chitinases.

### Acetate measurement using liquid-chromatography mass-spectrometry

To measure extracellular acetate in degrader monocultures, samples were taken throughout growth on colloidal chitin and centrifuged at 3’700 RPM for 10 minutes to pellet cells. The cell-free supernatant was removed and stored at -20°C until preparation for mass spectrometry.

To measure acetate, we used aniline derivatisation followed by liquid chromatography tandem mass spectrometry (LC-MS/MS). To prepare the samples, the pH was adjusted to 4.5 using 7.5 mM hydrochloric acid. Standard solutions containing acetate, lactate, propionate, butyrate and succinate were prepared with 8 serial dilutions starting from 1 mM. The derivatisation solution was prepared as follows (for 10 mL): 5 mL pure methanol, 4.6 mL LC MS grade water, 91 μL aniline (95% pure), 95.5g EDC, 200 μL hydrochloric acid 2M, 5 μL of internal standard solution (62 mg sodium formate-^13^C, 52 mg sodium acetate-^13^C_2_, 51 mg succinic acid-2,2,3,3-d4, 96 mg butyric acid-d8 in 1 mL LC MS grade water). The samples were derivatized at 4°C for 30 minutes and quenched with β-mercaptoethanol for a further 30 minutes at room temperature. Samples were diluted 1:50 in ddH_2_O prior to measurement.

All measurements were performed on a SCIEX 5500 QTRAP, coupled to an Agilent 1290 Infinity II autosampler and using a CORTECS UPLC C18, 90Å, 1.6 µm, 2.1 mm × 50 mm reversed phase column (Waters) with a guard column and 0.2 µm inline filter. The mobile phase consisted of 95% of buffer A (10 mM ammonium formate and 0.1% (v/v) formic acid in ddH_2_O) and 5% of buffer B (100% acetonitrile and 0.1% (v/v) formic acid in ddH_2_O) with a 5 minute isocratic gradient from 5% buffer A to 95% buffer A. Measurements were performed in positive mode using multiple reaction monitoring scans. Data were analysed using Skyline (Adams *et al*. 2020), where quality control and peak area calculations were performed for all standards and samples. Further analysis was performed in R (R Core Team 2021).

### Extracellular proteomics

Degrader monocultures for extracellular proteomics analyses were grown in 250 mL flasks with a culture volume of 25 mL until an OD_600_ of 1 or, if there was no growth on chitin, for 7 days. To harvest extracellular proteins, the cultures were centrifuged at 3’700 RPM for 15 minutes at 4°C to pellet the cells. The supernatants were then syringe-filtered through a 0.2 µm filter (Sarstedt) and 1 cOmplete™ EDTA-free Protease Inhibitor Cocktail mini tablet (Sigma Aldrich) was dissolved in each 25 mL sample. The samples were stored at -70°C until sample preparation and measurement.

For sample preparation, we used 0.5 mL of each supernatant to achieve a protein concentration ≤ 500 µg, as the most concentrated sample was around 500 µg. Reduction and alkylation of cysteine residues was achieved by adding DTT to a final concentration of 5 mM to each sample and incubating at room temperature for 20 minutes. Then, iodoacetamide was added to a final concentration of 15 mM to each sample, with a further 20 minutes incubation in the dark at room temperature. The iodoacetamide reaction was quenched by adding a further 5 mM of DTT.

Protein precipitation was carried out using a single-pot solid-phase enhanced protocol (Hughes *et al*. 2019). 100 µL SeraMag beads (Cytiva Life Sciences) were added to 2 mL Eppendorf tubes which were placed on a magnetic rack for 2 minutes. The bead buffer was then removed, and the beads were reconstituted in 500 µL ddH2O. The bead mixture was added to each sample, followed by 100% ethanol to achieve a final proportion of 50% (v/v) ethanol. The samples were mixed and then incubated at room temperature for 10 minutes on a linear rocker. Following incubation, the samples were placed back on the magnetic rack, and the supernatants were removed without disturbing the beads. 1.5 mL 80% ethanol was added to each sample and were mixed by vortexing. The supernatant was then removed after the samples were incubated on the magnetic rack for 2 minutes. This ethanol washing step was repeated for a total of three washes. After the last wash, 100 *µ*L of EPPS (200 mM, pH 8.5) was added to each sample, followed by Pierce Trypsin Protease (Thermo Fisher Scientific) to a 1:100 (w/w) final ratio. The samples were then sonicated in a water bath to reconstitute the beads and incubated overnight in a thermomixer (37°C, 1’000 RPM). The next day, samples were spun at 20’000 g for 1 minute to pellet the beads, then placed on the magnetic rack to allow the beads to settle. The supernatant containing peptides was recovered, vacuum-dried, and stored at -70°C until analysis.

Peptides were analysed online by LC-MS/MS. Reversed phase chromatography was performed using a Vanquish Neo UPLC system (Thermo Fisher Scientific) with a heated column compartment set to 50°C. Mobile Phase A consisted of 0.1% (v/v) formic acid in water, while Mobile Phase B was 80% (v/v) acetonitrile in water and 0.1% (v/v) formic acid. Approximately 1 μg of peptides were loaded onto a C18 analytical column (500 mm, 75 μm inner diameter), packed in-house with 1.8 μm ReproSil-Pur C18 beads (Dr. Maisch) fritted with Kasil, keeping constant pressure of 600 bar or a maximum flow rate of 1 μl/min. After sample loading, the chromatographic gradient was run at 0.3 μl/min and consisted of a ramp from 0% to 43% Mobile Phase B, followed by a wash with 100% Mobile Phase B, and a final re-equilibration step of 3 column volumes (total run time 90 minutes). Peptides from each sample were analysed on an Orbitrap HF-X mass spectrometer (Thermo Fisher Scientific) using an overlapping window data-independent analysis (DIA) pattern (Searle *et al*. 2018) consisting of a precursor scan, followed by DIA windows. Precursor scans were recorded over a 390–1’010 m/z window, using a resolution setting of 120’000, an automatic gain control (AGC) target of 1 × 106 and a maximum injection time of 60 ms. The RF of the ion funnel was set at 40% of maximum. A total of 150 DIA windows were quadrupole selected with an 8 m/z isolation window from 400.43 to 1’000.7 m/z and fragmented by higher energy collisional dissociation, HCD, (NCE = 30, AGC target of 1 × 106, maximum injection time 60 ms), with data recorded in centroid mode. Data were collected using a resolution setting of 15,000, a loop count of 75 and a default precursor charge state of +3. Peptides were introduced into the mass spectrometer through a 10 μm tapered pulled tip emitter (Fossil Ion Tech) via a custom nano-electrospray ionization source, supplied with a spray voltage of 1.6 kV. The instrument transfer capillary temperature was held at 275°C.

Raw files from the instrument were converted to .mzML format using MSConvert (Chambers *et al*. 2012) with 32-bit encoding precision and using SIM as spectra. Vendor-specific peak picking was used as the first filter and demultiplexing as the second filter to handle single MS1 scans and 150 DIA scans. The files were searched using DIA-NN (Demichev *et al*. 2020) with the .mzML files as input. Each degrader’s files were searched individually against the reference proteome obtained from NCBI **(Table 1)**. We used a precursor m/z range of 300-1’200, and a fragment ion m/z range of 200-2’000. We also used the ’match between runs’ option, as well as deep-learning based spectra, RTs and IMs prediction. A q value threshold of 0.01 was applied across all samples. For degrader 6D03, we were unable to map measured peptides to its reference proteome and any other degrader proteome or the common contaminants database cRAP (The Global Proteome Machine, n.d.). Further data analysis was carried out in R (R Core Team 2021). Proteins with more than 1 detected precursor in each sample were retained for analysis.

### Degrader enzyme incubation feeding experiment

Degrader ’digestates’ were prepared by growing each degrader in monoculture on colloidal chitin, then harvesting the cell-free supernatant by centrifuging the cultures for 15 minutes at 3’700 RPM and filtering thorough a 0.2 µm membrane. The supernatants were then incubated with colloidal chitin (2 g L^-1^) at a 1:1 ratio in 50 mL Falcon tubes for 7 days on a low-profile roller (IBI Scientific). After the incubation period, the tubes were centrifuged at 500 RCF to sediment the chitin, and the supernatant was stored at -20°C until its use in growth experiments.

For the digestate growth experiment, each degrader was precultured separately on Marine Broth 2216 and inoculated in monoculture on each of the digestates, including its own, in a flat-bottom 96 well plate. The digestates were mixed 3:1 with MBL without an additional carbon source. Growth was monitored using OD_600_ measurements in a plate reader (Tecan) set to 26.5°C, without shaking. As the data were not normally distributed (Shapiro test *p* < 0.05), Kruskal-Wallis and Dunn’s tests were used to assess differences in maximum OD_600_ between substrates for each degrader.

### Pairwise degrader interaction screen

Degraders were precultured individually on Marine Broth 2216 in 15 mL Greiner tubes and transferred to a colloidal chitin preculture for 3-7 days depending on the strain, so that all strains would be in the exponential phase on the same day. Following this chitin preculture, the strains were inoculated 1:1000 in colloidal chitin MBL in either mono- or cocultures. OD_600_ was measured daily, and samples for chitin quantification (Pontrelli and Sauer 2025) and qPCR (see section ’Strain-specific qPCR’) were taken daily.

Quantification of strain abundances with qPCR, and quantification of chitin, were performed for days 0 and 5 of the experiment. Fold changes in maximum culture OD_600_, lag time (time taken to reach OD_600_ > 0.1), and chitin remaining after 5 days of growth were calculated between monocultures and relevant cocultures. Strain abundances were used to calculate doubling times of individual strains in mono- and cocultures, which were used to categorize interactions as amensal (one strain reduces doublings compared to monoculture while other remains within 1 standard deviation of its monoculture), commensal (one strain remains comparable to monoculture while other increases doublings by more than 1 standard deviation), or exploitative (one strain reduces doublings while the other increases). Strain abundances were also compared directly to determine the ‘winner’ in each coculture (strain with the higher mean abundance, at least 1 standard deviation apart from the other strain’s mean abundance).

### Strain-specific qPCR

For qPCR, DNA was extracted using the DNAdvance kit (Beckman Coulter) according to the manufacturer’s instructions. Cultures were diluted to an OD_600_ between 0.1 and 0.3 prior to extraction. Strain-specific primers and probes were designed in Geneious and aligned against genomes of all other strains using BLAST to ensure that there would be no off-target binding **(Table 1)**. For qPCR, GoTaq Master Mix (Promega) was used. The recipe per reaction was 7.5 µL GoTaq Master Mix, 0.5 µL primers (premixed forward and reverse primers at a concentration of 30 µM each), and 1 µL water. All qPCR was carried out on a QuantStudio 3 System (Thermo Fisher), with the following thermocycling conditions: 95°C 2 min, (95°C 15 s, 60°C 1 min) x 40, 95°C 15 s, 60°C 1 min, 95°C 1 s.

Raw CT values were converted to ng µL^-1^ DNA using standard curves prepared for each strain. For this, DNA from a culture of each strain grown to OD_600_ 1 was extracted using the DNAdvance kit (Beckman Coulter), and the DNA concentration was measured using the Qubit Broad Range DNA kit (Thermo Fisher Scientific). Dilution series for each strain were made, and the CT values of each DNA concentration step were determined using qPCR. This allowed us to construct a calibration curve for each strain that was later used to convert raw CT values to ng µL^-1^ DNA, which we use as a proxy for the absolute abundance of a given strain.

### Inhibition experiments with 3D05

To test the effect of different initial conditions (starting ratios of degrader 3D05), degrader 6C06 and 6D03 were each inoculated with varying abundances of degrader 3D05, from 50% to 0.001%, with colloidal chitin MBL. Degrader abundance in the inoculum was calculated using values in **Table 1**.

To test whether delayed inoculation of degrader 3D05 could alleviate its inhibitory effect, degrader 6C06 and 6D03 monocultures and cocultures with 3D05 were all begun at the same time, and 3D05 was added to a subset of the monocultures after 1 or 3 days at 50% abundance relative to its co-culture partner.

To test whether 3D05 produces an inhibitory compound that enacts contact-independent effects on degraders 6C06, 6D03 and 3D05 were each grown on 20 mM GlcNAc MBL for 24 hours in 100 mL Erlenmeyer flasks. The supernatants were harvested by centrifugation at 4’500 RPM for 10 minutes, followed by filtration using a 0.2 µm syringe filter to generate cell-free supernatants. The supernatants were mixed with colloidal chitin MBL at a 1:1 ratio.

To test whether siderophore-mediated competition could be causing inhibition by degrader 3D05, degraders were grown in coculture with 3D05 with or without iron-supplemented colloidal chitin MBL. To produce iron-supplemented colloidal chitin MBL, a stock solution of iron(III) chloride was diluted to a final concentration of 50 µm, which we determined as an intermediate concentration at which to supplement iron in order to test whether siderophore-mediated competition is active in the coculture (Schiessl *et al*. 2017, Stubbendieck *et al*. 2019).

### Data deposition

The mass spectrometry proteomics data have been deposited to the ProteomeXchange Consortium via the PRIDE (Perez-Riverol *et al*. 2025) partner repository with the dataset identifier PXD079701, available using the reviewer login token eMckUQ5Vt9CD. The underlying data and code for all main and supplementary figures are available at https://codeberg.org/vhaavisto/chitin_degraders_manuscript.git.

## Results

### Enzymatic repertoires for chitin degradation and chitooligomer uptake

Chitin degraders are usually defined by their encoding and expression of hydrolytic chitinases in the CAZy families GH18 and GH19 (Beier and Bertilsson 2013). Accordingly, all seven degraders encode GH18 chitinases, and all but two also encode at least one GH19 chitinase **(Fig. 1A**. Chitinase copy numbers vary markedly between the degraders; most encode between two and four chitinases, while 6C06 encodes 10 and 6D03 encodes six **(Fig. 1A)**. Additionally, 6D03 encodes a chitinase with two distinct catalytic domains **(Fig. 1A)**. Although 6C06 has previously been reported to encode up to 13 chitinases (Enke *et al*. 2018, Pontrelli *et al*. 2022), we chose to focus on those specifically annotated as GH18 or GH19 chitinases by the latest version of dbCAN (Zheng *et al*. 2023) across all degraders.

**Figure 1:**
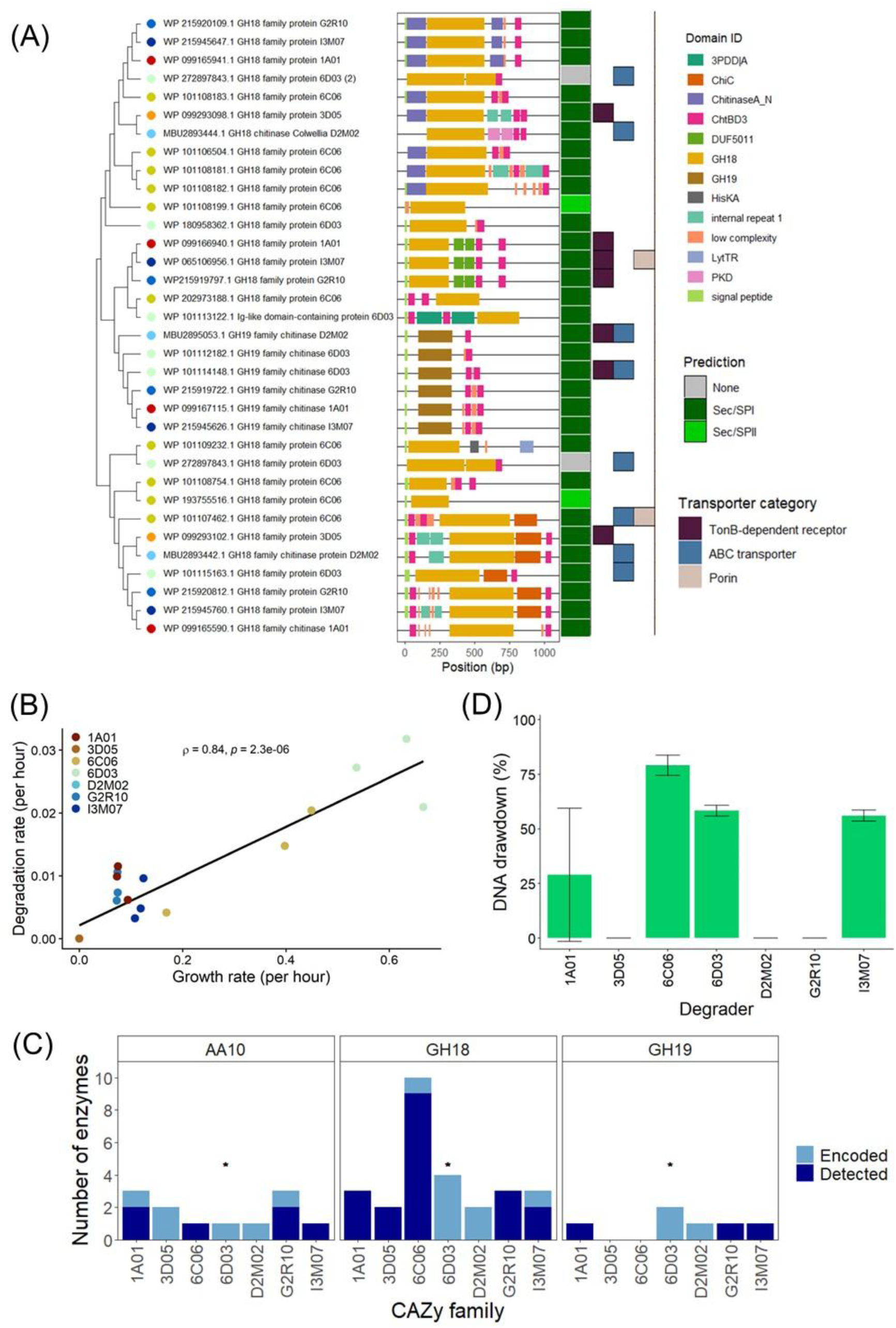
Genomic and physiological features of chitin degraders. **A)** Cladogram of GH18 and GH19 chitinases encoded by seven marine chitin degraders. The cladogram was constructed using aligned catalytic domain sequences in MEGA11. To the right of the cladogram are (from left to right) each enzyme’s domain architecture computed using EMBL SMART, predicted signal peptides computed with SignalP 6.0, and annotated transporters located within seven genes up- or downstream of each chitinase. **B)** Maximum growth and degradation rates of each degrader on colloidal chitin. Biological triplicates are shown separately. Spearman’s ρ and p-value for the correlation between growth and degradation rates are shown on the plot to the right of the legend. Growth and degradation rates were calculated using the Growthrates package in R. **C)** Attachment of chitin degraders to chitin particles. Attachment was measured using a draw-down assay, where cultures were incubated for 10 hours in minimal medium with or without 2 g L^-1^ colloidal chitin, and sampled from the planktonic phase. Total DNA was quantified using qPCR for each degrader **(Figure S6)**. Error bars indicate the standard deviation around the mean of three biological replicates. **D)** Genomic encoding and detection in proteomics measurements of selected CAZymes from all degraders. Asterisks indicate one degrader for which we were unable to quantify protein expression using proteomics. Abbreviations: CAZy, carbohydrate-active enzymes; GH18, glycosyl hydrolase family 18; GH19, glycosyl hydrolase family 19; CE4, polysaccharide/chitin deacetylase (carbohydrate esterase family 4); and AA10, lytic polysaccharide monooxygenases (auxiliary activity family 10).

In addition to determining chitinase copy number, we used EMBL SMART (Letunic and Bork 2018, Letunic, Khedkar, and Bork 2021) to investigate their domain architectures. Each degrader encodes several chitinases with diverse domain structures **(Fig. 1A)**, with 6C06 and I3M07 exhibiting the most structural diversity in terms of unique domains across their respective chitinases **(Fig. S1)**. Notably, every GH19 chitinase encoded by our degraders includes a predicted chitin-binding domain (ChtBD3), while many, albeit not all, GH18 chitinases include this domain. Many GH18 chitinases also contained either an N-terminal chitinase A domain (Pfam: ChitinaseA_N) or a C-terminal chitinase C domain (Pfam: ChiC), the functional roles of which remain uncertain (Perrakis, Ouzounis, and Wilson 1997).

The EMBL SMART annotations revealed that the majority of chitinases include an N-terminal signal peptide **(Fig. 1A)**, which targets proteins to the inner membrane for secretion into the periplasm (Kaushik, He, and Dalbey 2022). To better understand the localization of our chitinases, we used SignalP 6.0 (Teufel *et al*. 2022) to predict the types of signal peptides associated with each. The predictions suggest that most chitinases encode a Sec/SPI domain **(Fig. 1A)**, indicating that they are secreted into the periplasmic space via the Sec pathway (Kaushik, He, and Dalbey 2022). Two chitinases from 6C06 were predicted to encode a Sec/SPII domain, suggesting that the chitinase may undergo membrane anchoring after transport to the periplasm (Kaushik, He, and Dalbey 2022). Sec-domain-containing chitinases may be further transported from the periplasm to the extracellular space by type II or type V secretion systems (Green and Mecsas 2016), the latter of which are encoded by all degraders except 3D05 and D2M02 **(Fig. S2)**. One chitinase from 6D03 does not encode any predicted signal peptide, but could still be transported to the extracellular space directly by a Sec-independent process (Green and Mecsas 2016).

The genomic colocalization of chitinases and associated transporters may offer insights into the products generated by chitinases, as uptake mechanisms for chitooligomers should be tailored to match the products of specific chitinases (Liu *et al*. 2025). To investigate if chitin degraders employ such a strategy, we examined the genomic neighbourhood (spanning seven genes up- and downstream) of each chitinase. Our focus was on genes annotated as TonB-dependent transporters, porins, ABC transporters (excluding amino acid ABC transporters), and phosphotransferase systems, all of which are involved in chitooligomer transport (Hunt *et al*. 2008, Itoh and Kimoto 2019). Each degrader encodes at least one transporter-colocalized chitinase, although overall less than half of chitinases across all degraders were colocalized with a transporter **(Fig. 1A)**. Strains 3D05 and D2M02 had annotated transporters colocalized with all their chitinases, suggesting that they may optimize the uptake of degradation products specific to the activity of their respective enzymes more than the other strains. Together, these results indicate that the strategy of co-localizing transporters to chitinase products is present among our degraders, but not necessarily active across their entire chitinase repertoires.

### Growth and enzyme expression on colloidal chitin

Given the differences in size and diversity of our degraders’ chitinase repertoires, we hypothesized that particularly large (6C06) or structurally diverse (6C06, I3M07, 6D03) chitinase profiles might facilitate chitin degradation and robust growth. To assess this, we determined growth and degradation rates for each degrader on colloidal chitin and explored the relationship of these rates with copy number and structural diversity. We also assessed secretion of extracellular enzymes using proteomics.

In general, we observed a positive correlation between maximum growth and colloidal chitin degradation rates **(Fig. 1B)**, as well as a strong correlation between the number of chitinases and mean growth rate on colloidal chitin (Spearman’ rank correlation, ρ = 0.916, *p* = 0.004). No significant correlation was detected between chitinase structural diversity and growth rate. Degraders with the largest chitinase repertoires, specifically 6C06 and 6D03, also exhibited the highest growth and degradation rates **(Fig. 1B)**. Although 3D05 and D2M02 are classified as chitin degraders (Enke *et al*. 2018, Pollak *et al*. 2021) and encode chitinases **(Fig. 1A)**, they did not grow on colloidal chitin. The remaining strains showed similar, intermediate growth and degradation rates. For all degraders that grew, chitin degradation occurred within a 24-hour period, corresponding to the exponential growth phase **(Fig. S3)**.

To assess whether the entire chitinase repertoire is expressed during growth on colloidal chitin, we collected supernatants for extracellular proteomics from degrader monocultures that were grown on colloidal chitin, or, in cases where no growth was observed, after seven days of incubation **(Fig. S3)**. We analysed all proteins that were identified with at least two matching peptides **(Fig. S4)**, allowing for a false discovery rate of 1% and excluding measurements for degrader 6D03, where noise in the peptide measurements precluded peak identification and annotation. Most degraders showed detectable levels of all encoded GH18 and GH19 chitinases in the extracellular space **(Fig. 1C)**, with two exceptions: one GH18 chitinase from 6C06 and both GH18 chitinases from D2M02 were not detected. The lack of detection for D2M02 may be explained by its inability to grow on colloidal chitin, although both chitinases from 3D05 were detected despite its lack of growth. Overall, our recovery of nearly all encoded GH18 and GH19 chitinases suggests that our degraders deploy their full hydrolytic enzyme repertoires when growing on colloidal chitin.

In addition to the well-characterized hydrolytic pathway, chitin degradation may also proceed via oxidative reactions involving lytic polysaccharide monooxygenases (LPMOs) of the AA10 CAZy family, which encompassed chitin- and cellulose-active LPMOs (Vaaje-Kolstad *et al*. 2010). The oxidative pathway complements hydrolytic degradation by oxidising the surface of crystalline chitin, facilitating access for hydrolytic enzymes (Jiang *et al*. 2022). In particular, *Pseudoalteromonas* species are thought to utilize the oxidative pathway, although it has also been documented in *Vibrio* species (Jiang *et al*. 2022, Zhou *et al*. 2023). Every degrader in our study encodes at least one protein from the AA10 family, and four degraders (6C06, 1A01, I3M07 and G2R10) secreted AA10 family proteins during growth on colloidal chitin **(Fig. 1C)**. Although the normalized quantities of AA10 family proteins varied between and within degraders, many were present in quantities similar to the most abundant GH18 and GH19 chitinases in its extracellular proteome **(Fig. S5)**. This suggests a concurrent deployment of AA10 family LPMOs alongside hydrolytic chitinases for the degradation of colloidal chitin. However, since only one of the rapidly growing degraders expressed these enzymes, their precise role in promoting robust growth remains unclear.

Beyond enzyme encoding and expression, particle attachment may confer a competitive advantage during growth on chitin. Attached cells can efficiently take up degradation products near their source while minimizing diffusive losses, which may be consumed by others in the vicinity (Baty *et al*. 2000, Jagmann, Styp von Rekowski, and Philipp 2012, Frischkorn, Stojanovski, and Paranjpye 2013, Ebrahimi, Schwartzman, and Cordero 2019, Pontrelli *et al*. 2024). Alongside rapid growth and degradation, degraders 6C06 and 6D03 also demonstrated strong evidence of attachment to colloidal chitin **(Fig. 1D)**. This was evaluated using a draw-down assay that quantified the abundance of each strain in the planktonic phase of cultures incubated with or without colloidal chitin for 10 hours **(Fig. S6)**. 6C06 demonstrated the most pronounced attachment behaviour, achieving approximately 75% draw-down, suggesting that most of the cells in the population were attached to chitin particles, while both 6D03 and I3M07 showed around 55% draw-down. Given that two of the chitinases encoded by 6C06 may be bound to the outer membrane **(Fig. 1A)**, its attachment strategy may be linked to enzyme localization. The remaining degraders either failed to attach in this assay or yielded inconclusive results, suggesting that the three attaching degraders may be capable of privatizing degradation products, enhancing their ability to prevail in competitive interactions (Pontrelli *et al*. 2024).

### Low potential for degradation product niche partitioning among chitin degraders

Many of our degraders expressed both hydrolytic chitinases and LPMOs and exhibited robust growth on colloidal chitin. In particular, strains 6C06 and 6D03 displayed rapid growth rates and strong attachment to chitin particles, which may confer a competitive advantage in interactions. Since extracellular chitin degradation generates a shared resource pool composed of GlcNAc and chitooligomers of varying lengths, interaction outcomes will also depend on whether degraders compete for these resources, or partition them, for example according to chain length. Hydrolytic chitinases typically operate through two primary modes of action: exo-chitinases cleave diacetylchitobiose from the terminal ends of chitin chains, while endo-chitinases cleave randomly along the chitin chain, producing larger chitooligomers (Beier and Bertilsson 2013). Predicting degradation products based solely on chitinase sequences remains challenging (Guseva *et al*. 2024); however, *in vitro* enzyme assays have shown that 6D03 mainly produces GlcNAc, 6C06 mainly produces chitobiose, and strains 1A01, G2R10, and I3M07 produce a mix of chitooligomers ranging from one to four monomers in length (Pontrelli *et al*. 2022).

To investigate the potential for niche partitioning on colloidal chitin, we characterized the chitooligomer preferences of each degrader by measuring their maximum OD_600_ on GlcNAc and three degrader-derived chitooligomer mixes. These mixes were produced by incubating extracellular enzymes of 6D03, 6C06, and G2R10 with colloidal chitin *in vitro*, yielding mix G (predominantly GlcNAc), mix C (predominantly chitobiose), and mix GC (approximately equal parts of both), respectively (Pontrelli *et al*. 2022). To account for differences in total sugar content, we normalized maximum OD_600_ by the reducing sugar concentrations of each mix **(Fig. S7)** or the 20 mM GlcNAc control.

All degraders grew on the three mixes and on GlcNAc **(Fig. 2A)**, suggesting that they are generalist chitooligomer consumers. Mix GC (equal parts GlcNAc and chitobiose) yielded the highest maximum OD_600_ per mM of reducing sugars across all degraders **(Fig. 2A)**, indicating that a mix of chitooligomers was preferred compared to single chitooligomers; however, differences in maximum OD_600_ between the mixes were not statistically significant **(Table 2)**. Degraders 6D03, 6C06 and G2R10 also did not necessarily grow to higher OD_600_ on their own mixes compared to those derived from other degraders’ enzymes **(Fig S7)**, further suggesting that they consume a range of chitooligomers, with no clear preferences in terms of length.

**Figure 2:**
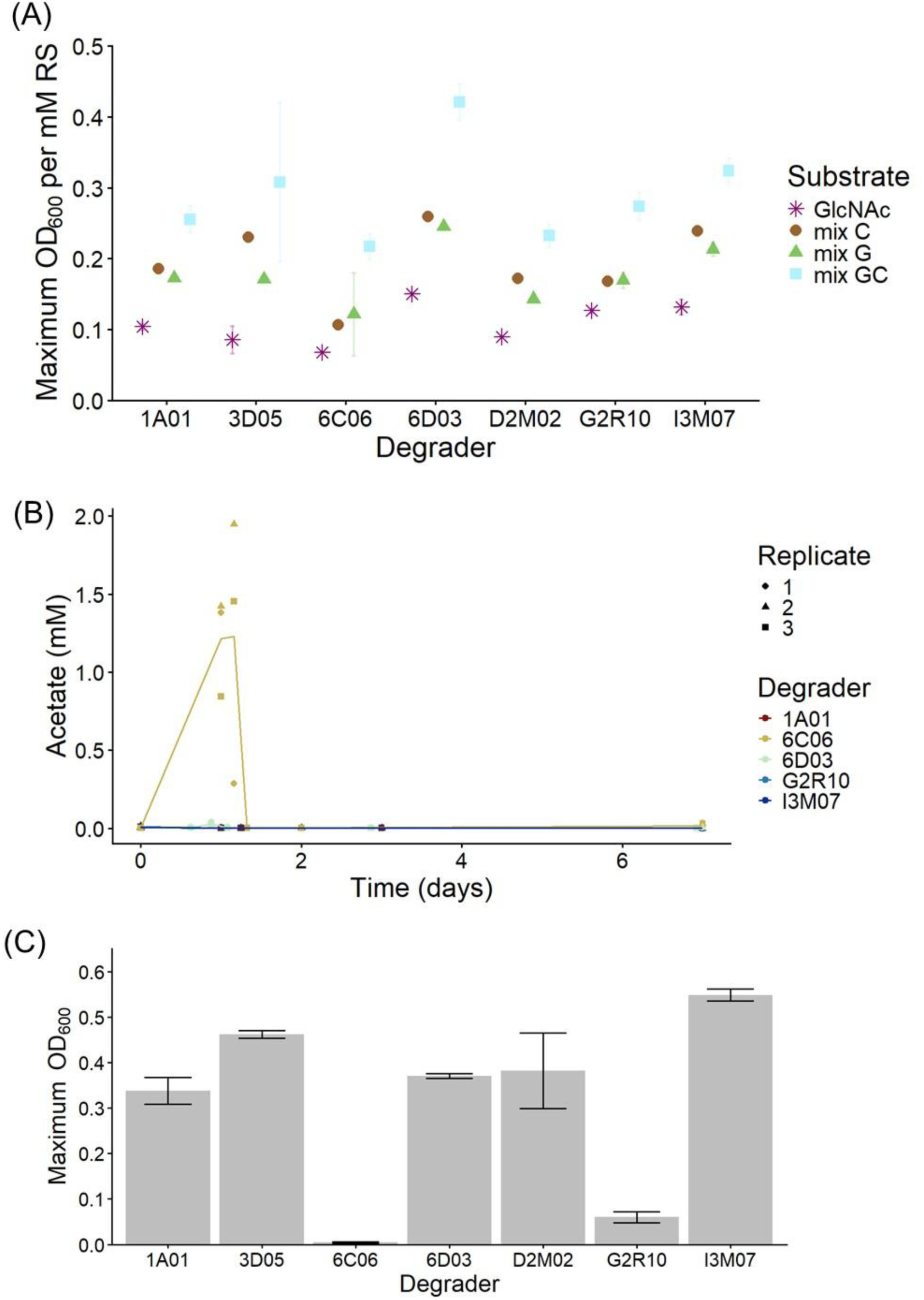
Opportunities for metabolic interactions and niche partitioning between degraders. **A)** Maximum OD_600_ of degraders on 20 mM GlcNAc and three chitooligomer mixes, normalized to the concentration of reducing sugars (RS) in each growth substrate **(Fig. S7)**. Mixes were obtained by incubating extracellular enzymes from 6C06, 6D03, and G2R10 with colloidal chitin. Mix G contains mainly GlcNAc, mix C contains mainly chitobiose, and mix GC consists of equal proportions of GlcNAc and chitobiose (Pontrelli *et al*. 2022). Kruskal-Wallis and Dunn’s tests were used to assess the significance of differences in maximum OD_600_ between the four substrates for each degrader **(Table 2)**. **B)** Acetate production over time for all degraders that grow in monoculture on colloidal chitin, measured using aniline derivatization followed by liquid chromatography-mass spectrometry. Biological replicates are shown separately by points, lines indicate the mean for each degrader. **C)** Growth of degraders on acetate. Maximum OD_600_ of degraders after five days of growth on 10 mM acetate as the sole carbon source. Error bars indicate standard deviation around the mean of three replicates.

**Table 2:** Dunn’s test results for maximum OD_600_ of degrader monocultures on chitooligomer mixes and GlcNAc. Significantly different comparisons (*p* adj. ≤ 0.05) are highlighted in bold.

| Degrader | Substrate1 | Substrate2 | n1 | n2 | Statistic | $p$ | $p$ adj. |
| --- | --- | --- | --- | --- | --- | --- | --- |
| 1A01 | GlcNAc | mix C | 3 | 3 | 2.038 | 0.042 | 0.083 |
| 1A01 | GlcNAc | mix G | 3 | 3 | 1.019 | 0.308 | 0.308 |
| **1A01** | **GlcNAc** | **mix GC** | **3** | **3** | **3.057** | **0.002** | **0.013** |
| 1A01 | mix C | mix G | 3 | 3 | -1.019 | 0.308 | 0.308 |
| 1A01 | mix C | mix GC | 3 | 3 | 1.019 | 0.308 | 0.308 |
| 1A01 | mix G | mix GC | 3 | 3 | 2.038 | 0.042 | 0.083 |
| D2M02 | GlcNAc | mix C | 3 | 2 | 1.628 | 0.103 | 0.207 |
| D2M02 | GlcNAc | mix G | 3 | 2 | 0.905 | 0.366 | 0.439 |
| **D2M02** | **GlcNAc** | **mix GC** | **3** | **3** | **2.832** | **0.005** | **0.028** |
| D2M02 | mix C | mix G | 2 | 2 | -0.661 | 0.509 | 0.509 |
| D2M02 | mix C | mix GC | 2 | 3 | 0.905 | 0.366 | 0.439 |
| D2M02 | mix G | mix GC | 2 | 3 | 1.628 | 0.103 | 0.207 |
| G2R10 | GlcNAc | mix C | 3 | 3 | 1.585 | 0.113 | 0.169 |
| G2R10 | GlcNAc | mix G | 3 | 3 | 1.472 | 0.141 | 0.169 |
| **G2R10** | **GlcNAc** | **mix GC** | **3** | **3** | **3.057** | **0.002** | **0.013** |
| G2R10 | mix C | mix G | 3 | 3 | -0.113 | 0.910 | 0.910 |
| G2R10 | mix C | mix GC | 3 | 3 | 1.472 | 0.141 | 0.169 |
| G2R10 | mix G | mix GC | 3 | 3 | 1.585 | 0.113 | 0.169 |
| I3M07 | GlcNAc | mix C | 3 | 3 | 2.038 | 0.042 | 0.083 |
| I3M07 | GlcNAc | mix G | 3 | 3 | 1.019 | 0.308 | 0.308 |
| **I3M07** | **GlcNAc** | **mix GC** | **3** | **3** | **3.057** | **0.002** | **0.013** |
| I3M07 | mix C | mix G | 3 | 3 | -1.019 | 0.308 | 0.308 |
| I3M07 | mix C | mix GC | 3 | 3 | 1.019 | 0.308 | 0.308 |
| I3M07 | mix G | mix GC | 3 | 3 | 2.038 | 0.042 | 0.083 |
| 6C06 | GlcNAc | mix C | 3 | 3 | 1.698 | 0.089 | 0.179 |
| 6C06 | GlcNAc | mix G | 3 | 3 | 1.359 | 0.174 | 0.209 |
| **6C06** | **GlcNAc** | **mix GC** | **3** | **3** | **3.057** | **0.002** | **0.013** |
| 6C06 | mix C | mix G | 3 | 3 | -0.340 | 0.734 | 0.734 |
| 6C06 | mix C | mix GC | 3 | 3 | 1.359 | 0.174 | 0.209 |
| 6C06 | mix G | mix GC | 3 | 3 | 1.698 | 0.089 | 0.179 |
| 6D03 | GlcNAc | mix C | 3 | 3 | 2.042 | 0.041 | 0.082 |
| 6D03 | GlcNAc | mix G | 3 | 3 | 1.021 | 0.307 | 0.307 |
| **6D03** | **GlcNAc** | **mix GC** | **3** | **3** | **3.063** | **0.002** | **0.013** |
| 6D03 | mix C | mix G | 3 | 3 | -1.021 | 0.307 | 0.307 |
| 6D03 | mix C | mix GC | 3 | 3 | 1.021 | 0.307 | 0.307 |
| 6D03 | mix G | mix GC | 3 | 3 | 2.042 | 0.041 | 0.082 |
| 3D05 | GlcNAc | mix C | 3 | 3 | 2.038 | 0.042 | 0.083 |
| 3D05 | GlcNAc | mix G | 3 | 3 | 1.019 | 0.308 | 0.308 |
| **3D05** | **GlcNAc** | **mix GC** | **3** | **3** | **3.057** | **0.002** | **0.013** |
| 3D05 | mix C | mix G | 3 | 3 | -1.019 | 0.308 | 0.308 |
| 3D05 | mix C | mix GC | 3 | 3 | 1.019 | 0.308 | 0.308 |
| 3D05 | mix G | mix GC | 3 | 3 | 2.038 | 0.042 | 0.083 |

In addition to chitooligomers, acetate can be produced during chitin degradation through extracellular deacetylation of chitin (Li X *et al*. 2007), cytoplasmic deacetylation of chitooligomers during conversion to fructose-6-phosphate (Bassler *et al*. 1991, Hunt *et al*. 2008, Jiang *et al*. 2022), or as a byproduct of overflow metabolism during rapid glycolytic growth (Paczia *et al*. 2012). Variations in acetate production and consumption capabilities among degraders could promote niche partitioning or intensify competitive interactions. Notably, only 6C06 exhibited transient accumulation of acetate in the extracellular space during exponential growth on colloidal chitin **(Fig. 2B)**. However, 6C06 was also the only degrader unable to grow on acetate as a sole carbon source **(Fig. 2C)**, suggesting that transiently accumulating extracellular acetate could be cross-fed in coculture settings with 6C06, possibly benefitting the partner degrader. Altogether, our findings do not provide evidence for niche partitioning chitooligomers among chitin degraders, but acetate could be crossfed in specific scenarios.

### Negative and neutral interactions prevail between degraders on colloidal chitin

As most degraders can utilize a broad range of chitin degradation products, we expected negative interactions to occur between them; these could manifest as competition for degradation products, or exploitation of the degradation products produced by another strain **(Fig. 3A**, (Ghoul and Mitri 2016)). Such negative interactions could result in reduced growth and lower chitin degradation in cocultures compared to monocultures. To test this hypothesis, we measured coculture growth, strain abundance, and residual chitin after five days of cultivation in mono- and cocultures of five degraders. Due to their rapid growth and attachment capabilities, we anticipated that 6D03 and 6C06 would outcompete slower-growing strains. Furthermore, we surmised that strains unable to grow on colloidal chitin in monoculture would exploit their coculture partners for degradation products, ultimately leading to reduced coculture growth and increased lag times (Pontrelli *et al*. 2026).

**Figure 3:**
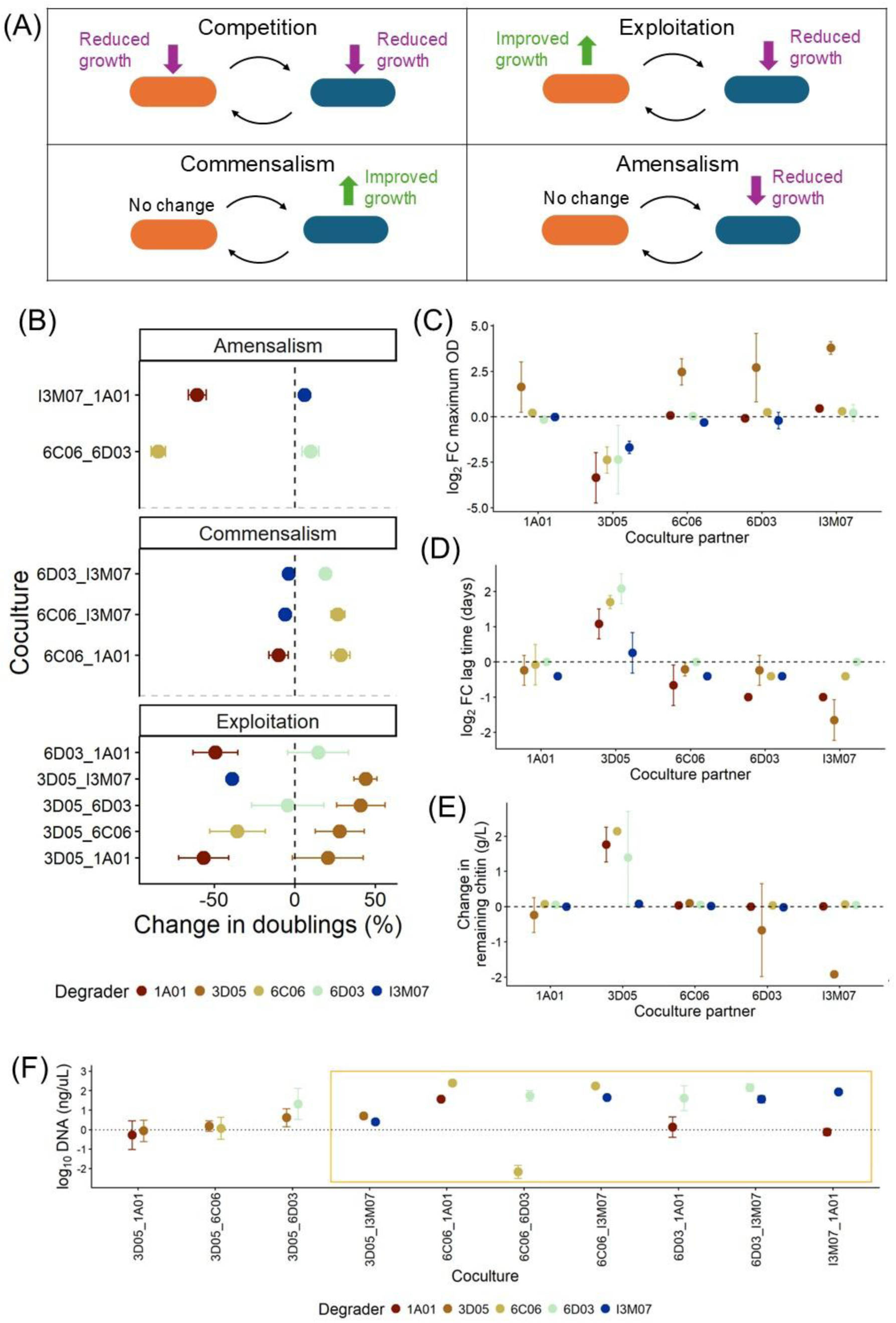
Pairwise interactions among degraders. **A)** Potential interactions between degraders and their impacts on interacting parties (adapted from (Ghoul and Mitri 2016)). **B)** Change in the number of doublings in each coculture pair relative to monocultures (dashed line). Differences in relative changes between the two strains of a given coculture were used to categorize interactions (facets, see Materials and Methods). **C)** Fold change in maximum OD_600_ of each degrader in coculture with a partner, relative to monocultures of the colour-coded strain. **D)** Fold change in coculture lag time of each degrader in coculture with a partner, relative to monocultures of the colour-coded strain. **E)** Change in quantity of chitin remaining after five days of growth for each degrader in coculture with a partner, relative to monocultures of the colour-coded strain. **F)** Strain abundances in coculture pairs. Abundance was determined by qPCR with strain-specific primers, using the amount of degrader DNA as a proxy for abundance. The orange box indicates cocultures where the difference between mean abundances of each degrader in each coculture exceeded 20%, indicating that the degrader with the higher abundance is the ‘winner’ in this interaction. All error bars represent the standard deviation around the mean of three biological replicates.

To determine the interaction type for each degrader pair **(Fig. 3A)**, we calculated the number of doublings for each strain in both mono- and cocultures using strain abundances on days 0 and 5. As expected, most interactions were negative; however, we did not detect any instances of competition, where both strains experience a reduction in doublings. Instead, we observed amensal interactions, in which one degrader was negatively affected while the other remained unaffected, and exploitative interactions where one degrader benefited at the expense of another **(Fig. 3B)**. Almost all exploitative interactions involved degrader 3D05, suggesting that it consistently exploits resources liberated by others, as expected given its lack of monoculture growth on colloidal chitin.

Despite mainly negative interactions among degraders, growth parameters such as maximum OD_600_ **(Fig. 3C)** and lag times **(Fig. 3D, S8),** as well as chitin degradation **(Fig. 3E)**, were largely comparable between mono- and cocultures. The recurrent exceptions were cocultures with 3D05, which exhibited lower maximum OD_600_, extended lag times, and decreased chitin degradation compared to the partner strain’s monoculture. Only the combination 3D05 and I3M07 did not show an increase in lag time or decreased chitin degradation **(Fig. 3D, E)**, despite reduced overall growth **(Fig. 3C)**, suggesting that I3M07 can overcome the initial exploitation-related growth delay.

We also detected commensalism **(Fig. 3A)** between three pairs of degraders, all including either 6C06 or 6D03 **(Fig. 3B)**, but this interaction did not improve coculture growth or chitin degradation despite the increased abundance of one degrader **(Fig. 3C-E)**. Indeed, none of the cocultures exhibited synergy, defined as growth greater than the sum of both members’ monocultures **(Fig. S9)**, indicating that interactions between degraders did not result in cooperative degradation.

To identify which strain was more successful in each coculture, we assessed abundances after five days of growth. Consistent with their robust growth and attachment behaviour, 6C06 and 6D03 proved to be the most successful strains, surpassing their coculture partners in two and three out of four cases, respectively **(Fig. 3F)**. Despite its exploitative behaviour, 3D05 did not outcompete its coculture partner in most cocultures **(Fig. 3F)**. Together with the large reductions in chitin degradation in most cocultures with 3D05, these results demonstrate that the exploitative dynamic is severe enough to substantially curtail chitin degradation, with 3D05 failing to make a profit from its exploitation of other degraders.

### Degrader 3D05 may inhibit others through secreted compounds

How 3D05 inhibits other degraders remains an unanswered question; particularly surprising was the strong inhibitory effect on 6D03, which grew rapidly on both colloidal chitin and GlcNAc in monoculture **(Fig. 1B, 2A)**. Unlike 3D05, 6D03 and 6C06 also both attach to colloidal chitin, which could help them to privatise chitooligomers and reduce the effectiveness of exploitation (Pontrelli *et al*. 2026) **(Fig. 2C)**. However, I3M07 also attaches to chitin but was not strongly inhibited by 3D05 **(Fig. 3C, 3D)**, suggesting that attachment is not the only mechanism determining the outcome of these interactions.

Certain non-degrading ‘exploiter’ strains can completely inhibit the growth of chitin degraders in a manner comparable to what we observed between 3D05 and 6D03, by siphoning GlcNAc in the early stages of culture growth (Pontrelli *et al*. 2026). This inhibitory effect is eliminated when the exploiter strain is seeded at densities 20-fold lower than its partner degrader (Pontrelli *et al*. 2026). To test whether this inhibitory mechanism is also employed by 3D05, we co-inoculated the two most successful degraders, 6D03 and 6C06, with 3D05 at different starting densities relative to its coculture partner and measured culture lag time, which can serve as an indicator for inhibitory dynamics (Pontrelli *et al*. 2026). The inhibitory effect persisted across a wide range of 3D05 starting densities for both partner degraders, with no significant relationship between 3D05 abundance and lag time for 6D03 (Spearman’s rank correlation, ρ = 0.41, *p* = 27) or 6C06 (Spearman’s rank correlation, ρ = 0.47, *p* = 0.05). As the effect on 6D03 remained strong at 1% 3D05, we tried inoculating at an additional three concentrations compared to 6C06. However, even inoculating 3D05 at 0.001% compared to 6D03 did not reproducibly restore coculture growth **(Fig. 4A)**. We concluded that the inhibitory mechanism of 3D05 is probably unrelated to initial siphoning of GlcNAc and sought out another explanation for the inhibition we observed.

**Figure 4:**
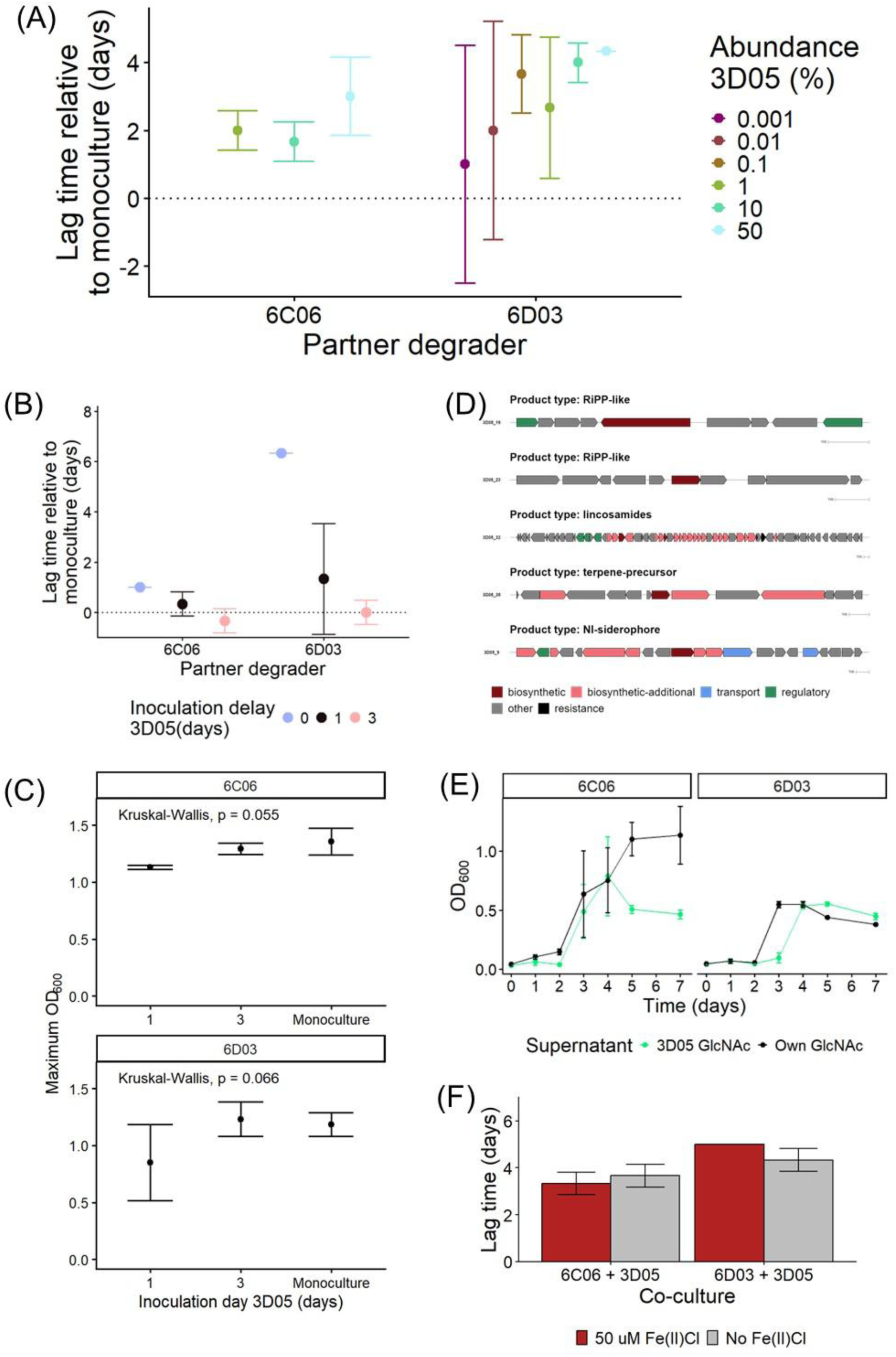
Investigating the mechanism of inhibition by 3D05. **A)** Effect of changing the inoculation ratio of 3D05 on growth of cocultures. Lag times are shown relative to the monoculture of either 6D03 or 6C06 (dotted lines). The abundance of 3D05 is expressed relative to that of its partner degrader. **B)** Effect of delaying inoculation of 3D05 on coculture lag time with 6C06 or 6D03. Lag times are shown relative to the monoculture of either 6D03 or 6C06 (dotted lines). **C)** Effect of delaying inoculation of 3D05 on maximum OD_600_ in coculture with 6D03 or 6C06. Kruskal-Wallis tests were performed to compare the maximum OD_600_ across inoculation conditions for each degrader, results are reported on the plots. **D)** Five predicted biosynthetic gene clusters encoded by 3D05. Prediction was done using antiSMASH v8, predicted gene functions within each cluster are colour-coded. **E)** Growth of 6D03 and 6C06 with supplementation of supernatant from their own monoculture or 3D05 monocultures. Supernatants were generated by growing each strain on 20 mM GlcNAc for 24 hours and passing these cultures through a 0.2 µm filter to remove cells. Supernatants were supplemented at a 1:1 concentration with colloidal chitin MBL. **F)** Lag time of cocultures with 3D05 and 6C06 or 6D03, with or without iron(II) chloride supplementation, in addition to the trace metals mix normally supplied in growth media (see Materials and Methods). All error bars indicate standard deviation around the mean of three biological replicates.

We next tested whether delaying the inoculation of 3D05 could alleviate the inhibitory effect, by allowing its partner degrader to begin chitin degradation in its absence. For 6D03, delaying 3D05 inoculation by one day was sufficient to restore growth comparable to the 6D03 monoculture, while for 6C06, delaying by three days restored monoculture-like growth **(Fig. 4B)**. These results suggest that initial conditions are important, with a given threshold of the partner degrader being required to avoid inhibition by 3D05. As the maximum OD_600_ of cocultures was not affected following delayed inoculation **(Fig. 4C)**, we could rule out contact-dependent inhibition as a mechanism. Contact-dependent inhibition through type V or VI secretion systems should be effective even when 3D05 is at a relatively low abundance compared to its target (Booth, Smith, and Foster 2023), and 3D05 also does not encode a type V or VI secretion system. Thus, we are left to consider further inhibitory mechanisms that could be sensitive to initial conditions.

Marine *Pseudoalteromonas* species such as 3D05 are known to be prolific producers of secondary metabolites including antibiotics and siderophores, as well as proteases (Chen XL *et al*. 2020, Wang *et al*. 2024). Indeed, 3D05 is predicted to encode five biosynthetic gene clusters encoding for compounds including a lincosamide antibiotic and a nickel-iron siderophore **(Fig. 4D)**. To test whether 3D05 produces compounds that can inhibit other degraders, we grew 3D05, 6C06 and 6D03 in monoculture on 20 mM GlcNAc, and supplemented supernatants from these monocultures to 6D03 and 6C06. 6D03 experienced an increased lag time when exposed to 3D05 supernatant compared to its own supernatant **(Fig. 4E)**. For 6C06, lag time did not differ when grown on 3D05 supernatant compared to its own, although we observed a decrease in final OD_600_ with 3D05 supernatant **(Fig. 4E)**. In both cases, we concluded that 3D05 produces a compound that can attenuate the growth of others, even in the absence of other strains in its environment.

Secondary metabolite expression in bacteria can be modulated by external conditions, including environmental factors and the presence of other strains (Gasparek, Steel, and Papachristodoulou 2023). One secondary metabolite that could be expressed in monoculture conditions is the predicted nickel-iron siderophore **(Fig. 4D)**. Iron competition using siderophores is thought to be most prevalent in the early stages of growth and may therefore be sensitive to initial conditions (Leinweber, Fredrik Inglis, and Kümmerli 2017). This mechanism could allow 3D05 to siphon iron from its competitors if they lack the appropriate siderophore receptor (Schiessl *et al*. 2017). To investigate whether competition for iron drives the inhibitory effect of 3D05, we grew it in coculture with 6C06 and 6D03 with or without the addition of 50 µM iron(III) chloride, to alleviate any iron competition by providing it in excess. Our minimal marine medium normally contains 12.9 µM iron(III) chloride, which can be considered an intermediate concentration (Schiessl *et al*. 2017). We observed no impact of iron supplementation on the lag time in cocultures of 6C06 or 6D03 with 3D05 **(Fig. 4F)**, indicating that siderophore-mediated competition is unlikely to be active. Thus, the remaining products of the predicted biosynthetic gene clusters remain as likely drivers of the inhibitory effect.

## Discussion

We demonstrate that taxonomically diverse marine chitin degraders exhibit generalist resource use behaviours, consuming a range of chitooligomers. Although we expected these consumption patterns to lead to competition, exploitation, amensalism and commensalism were prevalent between degraders growing on colloidal chitin. Interestingly, only exploitative interactions led to reduced chitin degradation, despite changes in the growth of degraders in all three types of interactions. Negative interactions such as exploitation have only been reported in one pair of chitin degraders (Jagmann, Styp von Rekowski, and Philipp 2012), but are well-known from interactions between chitin degraders and non-degraders (Daniels *et al*. 2022, Pontrelli *et al*. 2026). Our work suggests that negative interactions are relatively frequent among marine chitin degraders, though they do not always lead to reduced degradation.

In contrast to work with polysaccharides such as xylan (Abdoli *et al*. 2024) and the brown algal polysaccharide fucoidan (Sichert *et al*. 2025), we did not detect synergistic degradation between multiple degraders. This suggests that the challenges associated with utilizing homopolysaccharides such as chitin – structures composed of a single monomer type linked by a single bond type – may differ from those associated with utilizing heteropolysaccharides such as xylan and fucoidan, which contain two or more distinct monomers and different linkage types. Indeed, negative interactions have also been reported between degraders of other homopolymers such as cellulose, where different taxa can competitively exclude each other through mechanisms such as substrate attachment, superior degradation product affinities, and production of inhibitory compounds (Shi, Odt, and Weimer 1997, Chen J and Weimer 2001, Wilhelm *et al*. 2021, Lewin *et al*. 2022).

Interestingly, all degraders expressed near-complete GH18 and GH19 chitinase repertoires during growth on colloidal chitin, with some degraders also expressing AA10 family enzymes associated with oxidative chitin degradation (Jiang *et al*. 2022). The reason for a single degrader to encode multiple different degrading enzymes for a single polysaccharide remains an open question; while some complex polysaccharides require concerted action of many different enzymes (Sichert *et al*. 2020), chitin degradation can in principle be achieved with a single hydrolytic enzyme (Beier and Bertilsson 2013). Previous work has shown that *Vibrio* species express some chitinases during growth on various types of chitin, but also specifically express certain chitinases in response to specific chitin types (Svitil *et al*. 1997). However, chitinase expression did not always translate to growth in our experiments, as two degraders did not grow on colloidal chitin despite producing detectable chitinases. Degrader 3D05 has previously been cultivated in monoculture on chitin beads (Enke *et al*. 2018), whereas D2M02 lacks documented growth on chitin in monoculture despite being categorized as a chitin degrader (Pollak *et al*. 2021). Chitin degradation has also not been reported in other members of the genus *Colwellia*, despite the presence of chitinases in some strains’ genomes (Raimundo *et al*. 2021).

Large chitinase repertoires and the ability to attach to chitin particles, seen for degraders 6C06 and 6D03, may contribute to success in cocultures. However, accurately predicting interaction outcomes also requires consideration of inhibitory effects resulting from secreted compounds (Schmitz *et al*. 2024), as we saw in cases where fast-growing degraders were strongly inhibited by degrader 3D05. This finding raises questions about the strategies chitin degraders employ to gain a competitive advantage. The efficacy of inhibition during the initial stages of degradation implies that degrader 3D05 may deter competitors from accessing nutrients before degradation can be initiated, ultimately stalling growth altogether. Given the exclusion of alternative mechanisms such as exploitation of chitin degradation products or siderophore-based nickel-iron scavenging, the most likely explanation is secretion of an inhibitory compound. Degrader 3D05 is predicted to encode five biosynthetic gene clusters, including one for a lincosamide antibiotic and two for ribosomally synthesized and post-translationally modified peptide (RiPP)-like products. The structures and bioactivities of the RiPP-like products are unknown; however, RiPPs have been implicated in microbial competition through mechanisms such as pore formation in target cells and modulation of quorum sensing responses (Li Y and Rebuffat 2020). Lincosamide production has not previously been reported in *Pseudoalteromonas* or other marine bacteria, as this class of antibiotics is predominantly associated with soil-dwelling *Streptomyces* species (Mori and Abe 2024).

Community assembly on chitin begins with degrader colonisation and initiation of degradation, and is followed by the recruitment of various non-degrading strains through the generation of a public resource pool (Datta *et al*. 2016, Enke *et al*. 2019, Pontrelli *et al*. 2022). This process has been studied in detail for communities of single degraders and multiple non-degraders (Pontrelli *et al*. 2022); however, the collective effects of multiple degraders on community assembly have not yet been examined. The presence of several degraders may lead to the production and liberation of a greater quantity or a more diverse array of metabolites into the public resource pool, which could establish additional niches for non-degrading strains. Furthermore, interactions among degraders could be influenced by the introduction of non-degrading strains, for example through higher-order interactions that can reduce the effects of negative interactions (Sundarraman *et al*. 2020). Our findings show that certain degrader combinations are more productive than others, but whether interactions proceed similarly in the presence of a more complex community remains an open question.

## Acknowledgements

We thank Kian Bigović Villi for help with genomic analyses. We also gratefully acknowledge the Simons Foundation Principles of Microbial Ecosystems (PriME) collaboration for funding, and thank PriME members for providing valuable feedback at various stages of the project.

## Author contributions

Vilhelmiina Haavisto (Conceptualization, Investigation, Formal analysis, Data curation, Visualization, Writing – original draft, Writing – review & editing), Peter F. Doubleday (Methodology, Investigation, Resources), Andreas Sichert (Conceptualization, Supervision, Writing – review & editing, Methodology, Resources), Uwe Sauer (Conceptualization, Writing – review & editing, Supervision, Funding acquisition)

