## Supplementary Materials for "Negative and neutral interactions are prevalent in interactions between marine chitin degraders"

Includes Supplementary Figures S1-S9

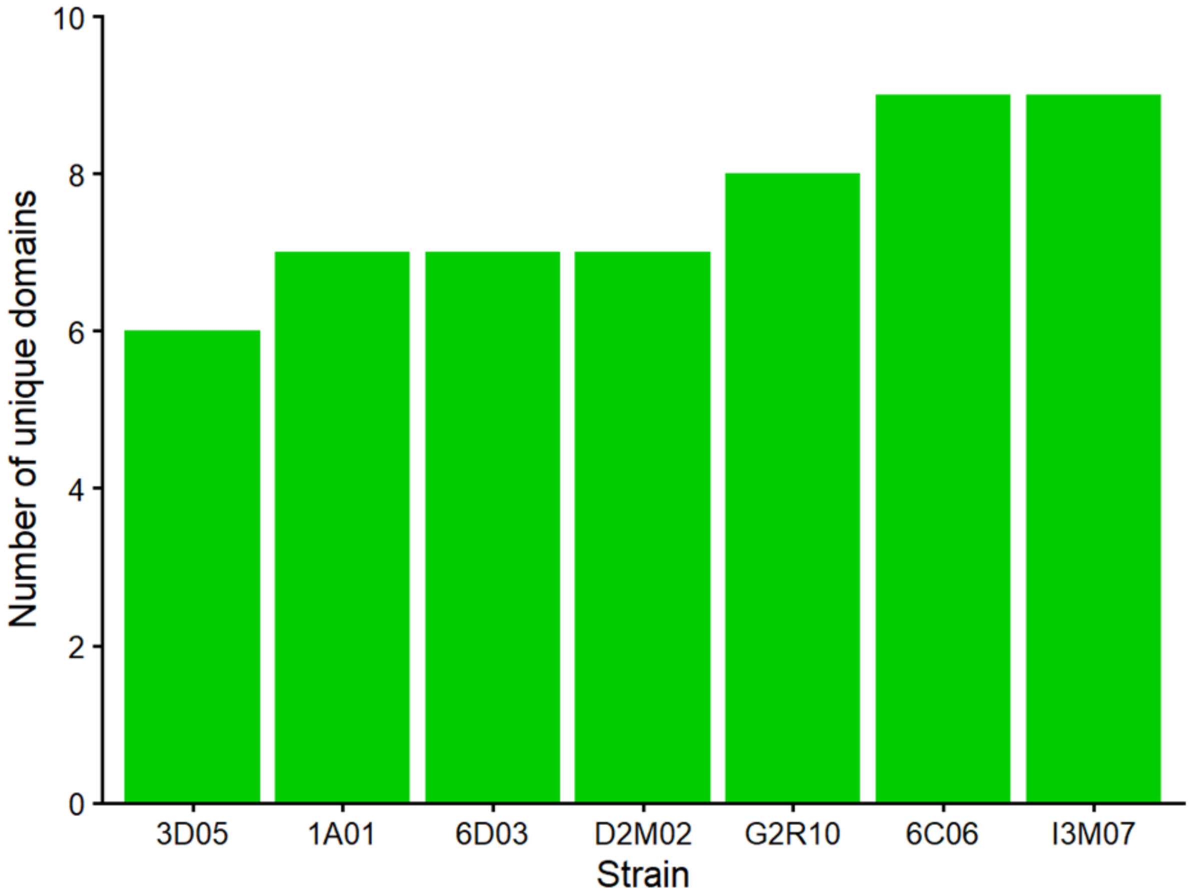


**Supplementary Figure S1:** Number of unique domains in each degrader's GH18 and GH19 chitinases predicted using EMBL SMART.

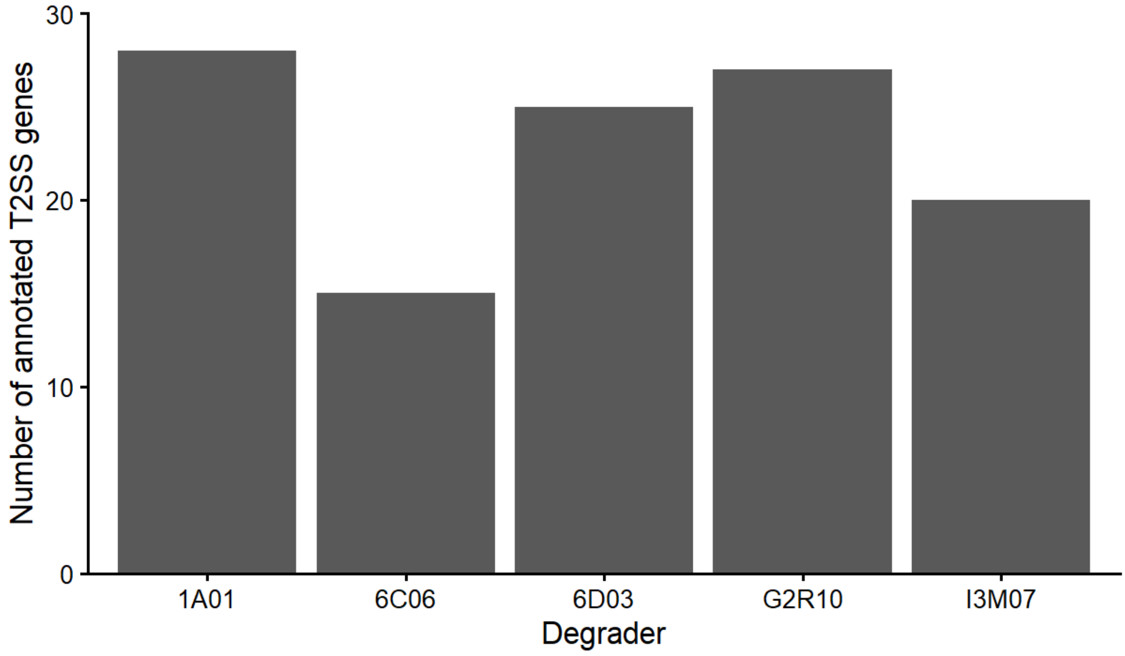


**Supplementary Figure S2**: Encoding of type II secretion system (T2SS) genes by degraders. Annotated genomes were searched for the terms “type II secretion system” and “type V secretion system”, the latter of which produced no hits. Degraders 3D05 and D2M02 did not have any annotated genes with either search term and are not shown in the figure.

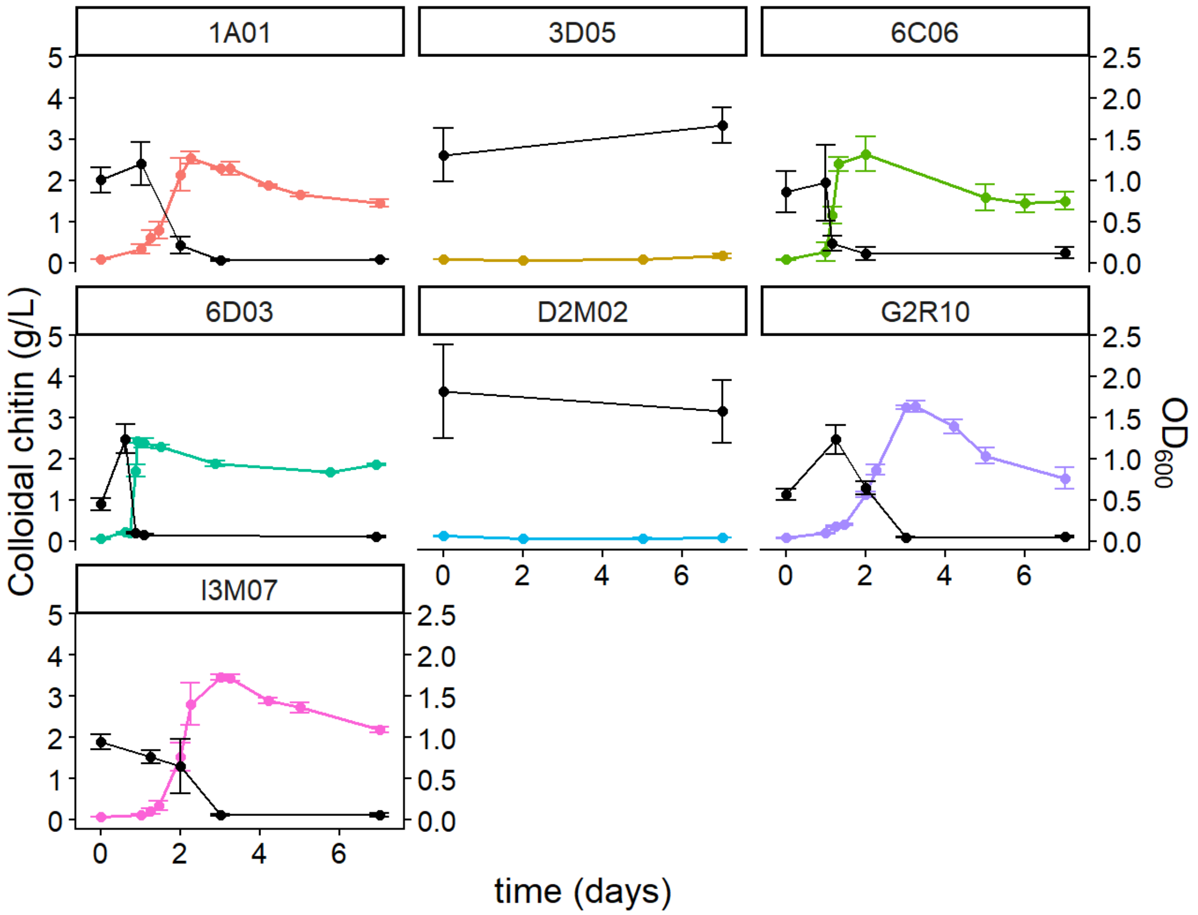


**Supplementary Figure S3**: OD600 growth curves of all degraders on colloidal chitin (colours, right axis) alongside chitin degradation (black lines, left axis). Error bars show the standard deviation around the mean of three replicates.

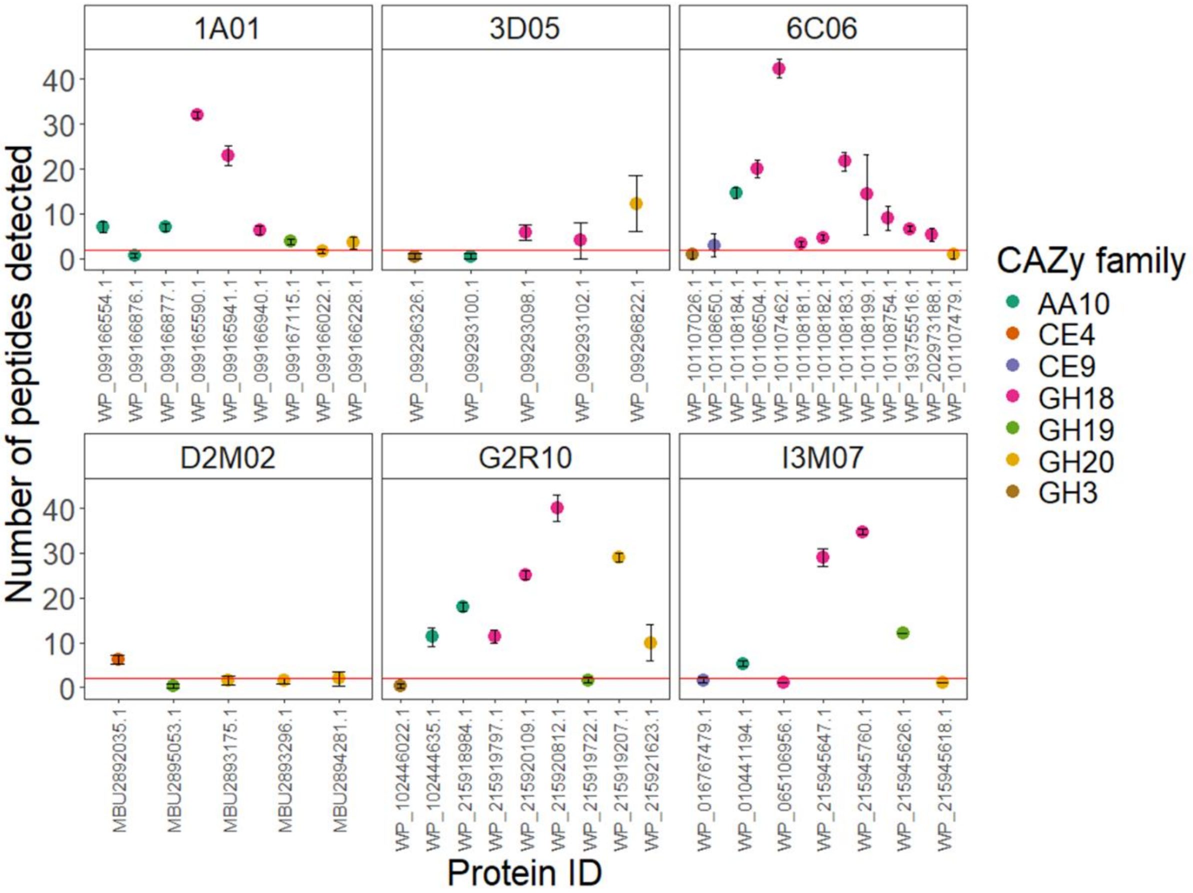


**Supplementary Figure S4:** Number of precursor peptides detected for each protein included in analysis of extracellular enzyme expression in degraders growing in monoculture on colloidal chitin. The red line indicates the threshold at which proteins were considered "expressed" (minimum 2 precursors detected). Error bars indicate the standard deviation around the mean of two biological replicates.

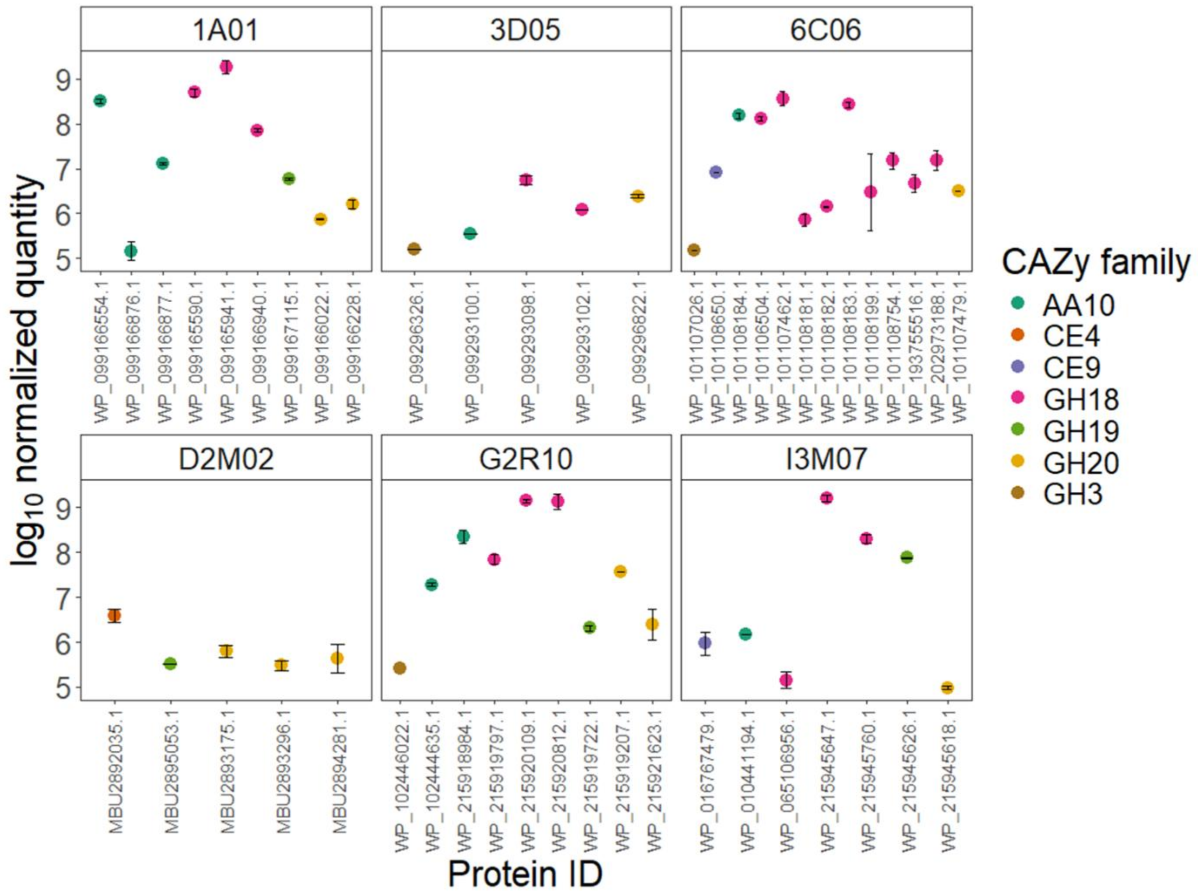


**Supplementary Figure S5:** Normalized quantities of precursors detected for each protein included in analysis of extracellular enzyme expression in degraders growing in monoculture on colloidal chitin. Error bars indicate the standard deviation around the mean of two biological replicates.

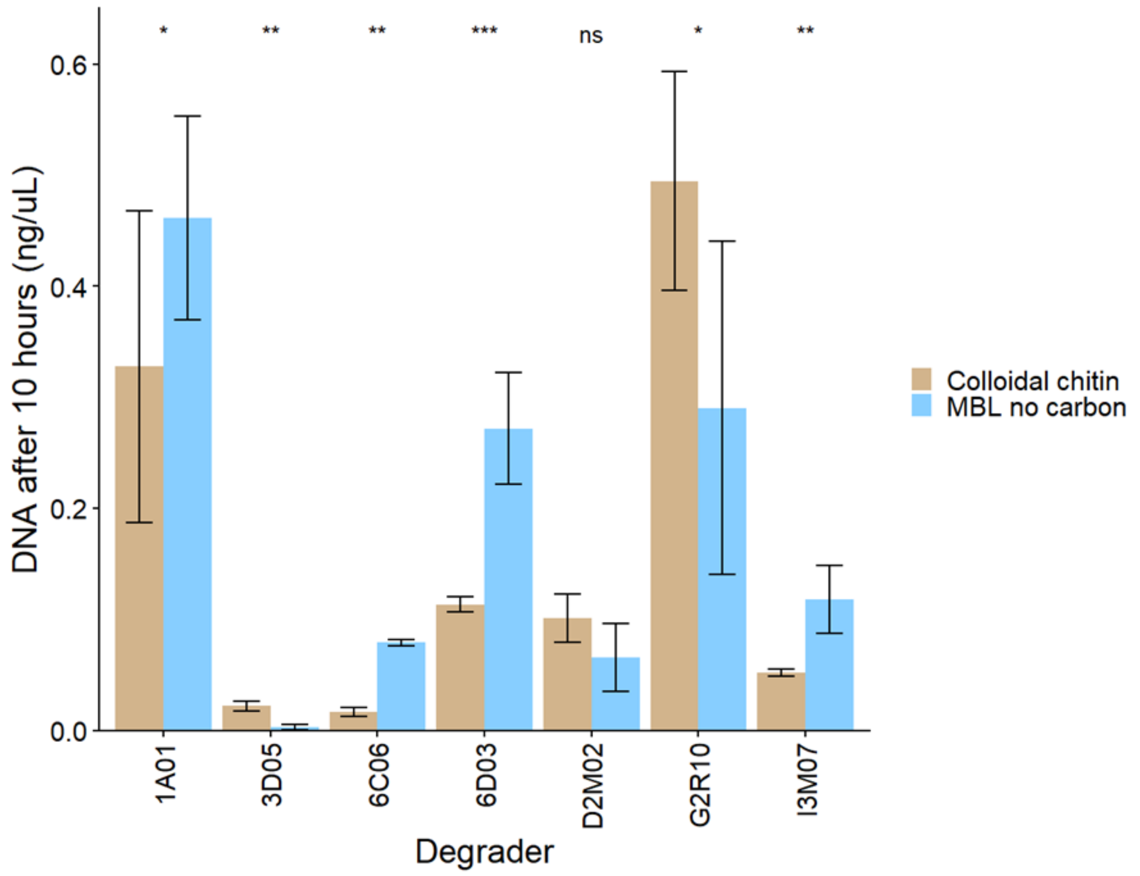


**Supplementary Figure S6**: DNA concentration for all degraders in cultures incubated with or without 2 g $L^{-1}$ colloidal chitin for 10 hours. Asterisks indicate significant differences between DNA concentrations in each condition for each degrader, measured using a t-test (*** = p<0.0001. ** = p<0.001, * = p<0.01, ns = not significant (p>0.5)).

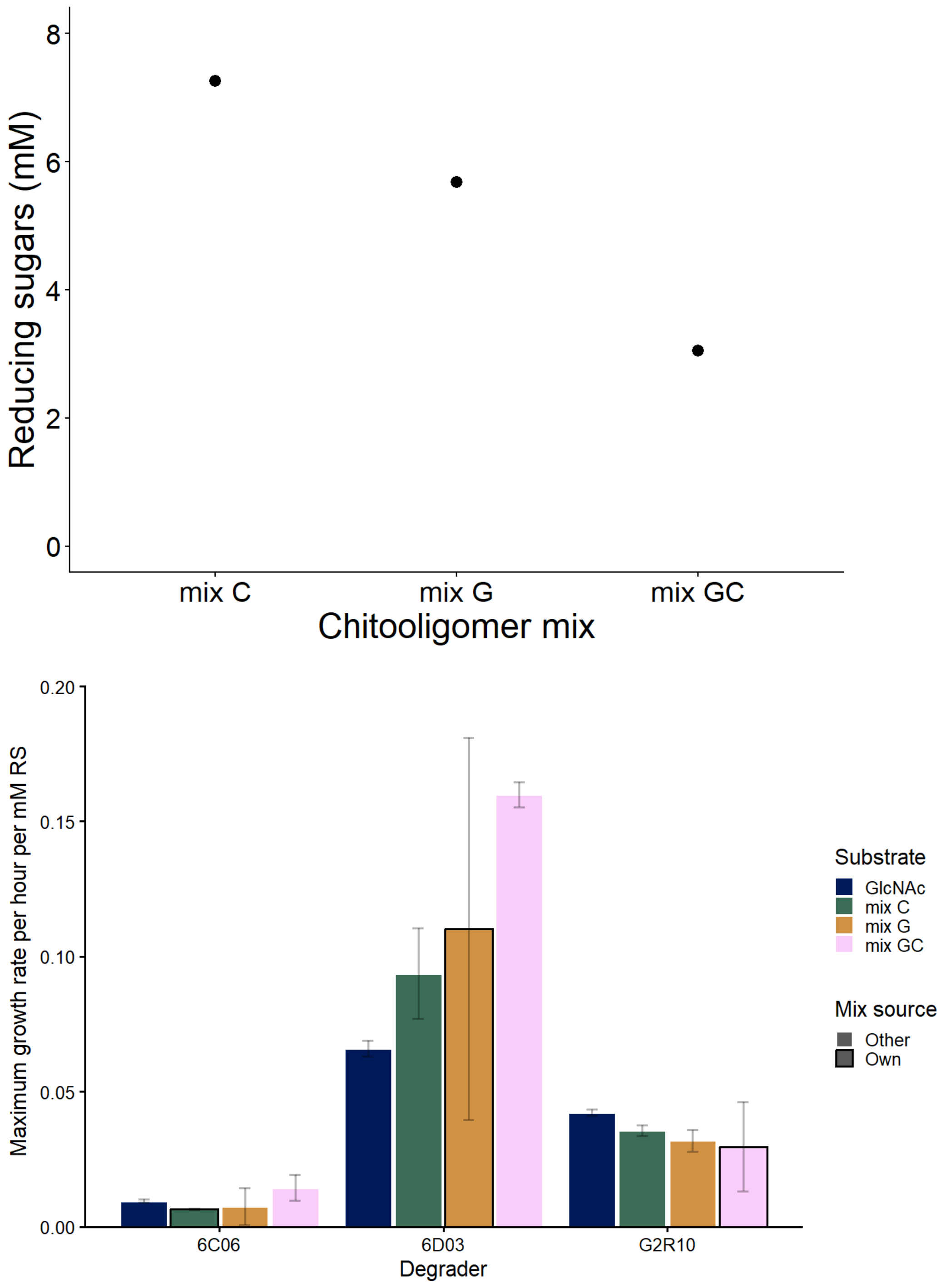


**Supplementary Figure S7**: Reducing sugar content of chitooligomer mixes derived from degrader enzymes incubated with colloidal chitin, and growth of degraders from which the chitooligomer mixes were derived on all three mixes and 20 mM GlcNAc. Error bars show standard deviation around the mean of two technical replicates (upper plot) and three biological replicates (lower plot).

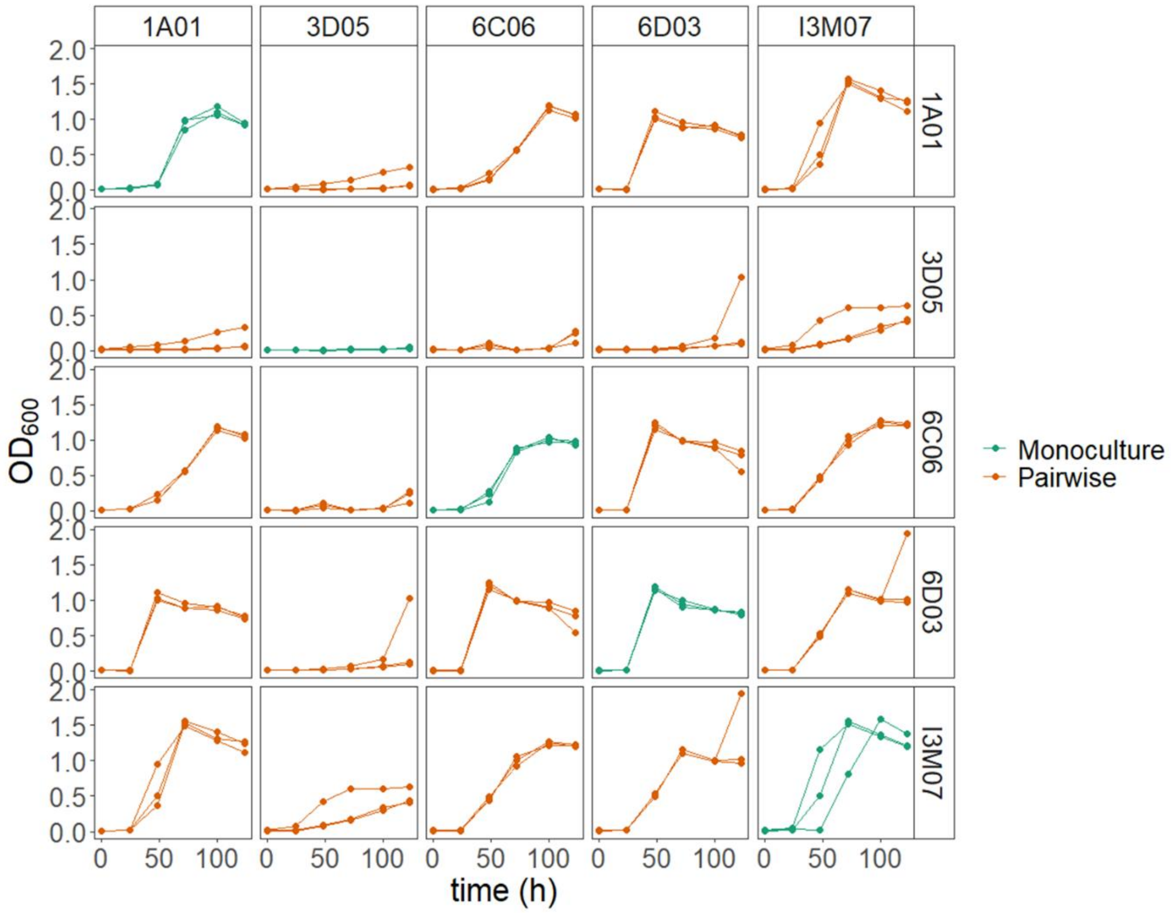


**Supplementary Figure S8:** $OD_{600}$ growth curves of all degrader monocultures and pairwise cocultures on 2 g $L^{-1}$ colloidal chitin. Biological replicates are shown separately.

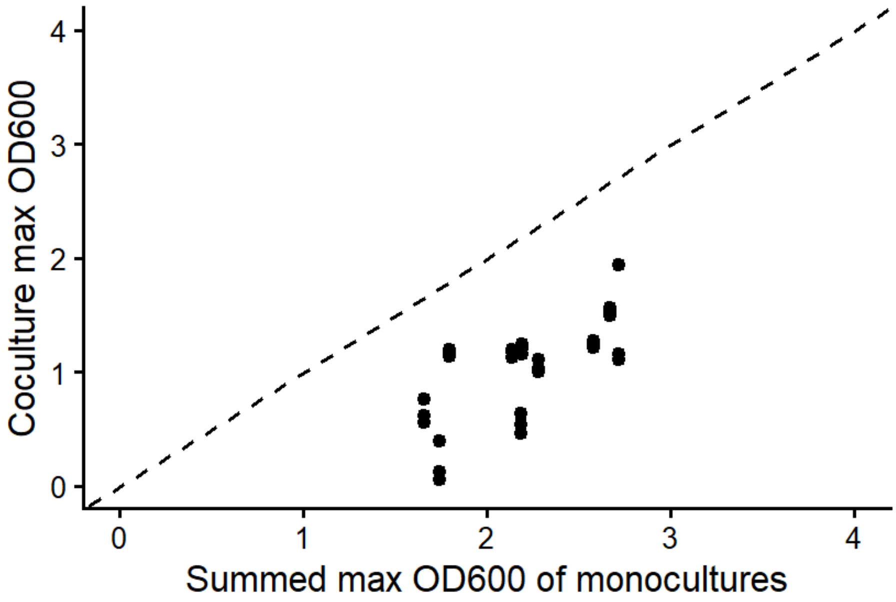


**Supplementary Figure S9:** Summed maximum $OD_{600}$ of degrader monocultures plotted against the maximum $OD_{600}$ of pairwise degrader cocultures. The dotted line indicates a theoretical 1:1 relationship.
